# Genome-resolved metagenomics reveals abundant nitrate reducers and partitioning of nitrite usage within global oxygen deficient zones

**DOI:** 10.1101/2023.03.02.530666

**Authors:** Irene H. Zhang, Xin Sun, Amal Jayakumar, Samantha G. Fortin, Bess B. Ward, Andrew R. Babbin

**Affiliations:** Department of Earth, Atmospheric and Planetary Sciences, Massachusetts Institute of Technology, Cambridge, MA; Program in Microbiology, Massachusetts Institute of Technology, Cambridge, MA; Carnegie Institution for Science, Department of Global Ecology, Stanford, CA; Department of Geosciences, Princeton University, Princeton, NJ

## Abstract

Oxygen deficient zones (ODZs) account for about 30% of total oceanic fixed nitrogen loss via processes including denitrification, a microbially-mediated pathway proceeding stepwise from NO_3_^−^ to N_2_. This process may be performed entirely by complete denitrifiers capable of all four steps, but many organisms possess only partial denitrification pathways, either producing or consuming key intermediates such as the greenhouse gas N_2_O. Marker gene surveys have revealed a diversity of denitrification genes within ODZs, but whether these genes are primarily carried by complete or partial denitrifiers and the identities of denitrifying taxa remain open questions. From 56 metagenomes spanning all three major ODZs, we use genome-resolved metagenomics to reveal the predominance of partial denitrifiers, particularly single-step denitrifiers. We find niche differentiation among nitrogen-cycling organisms, with communities performing each nitrogen transformation distinct in taxonomic identity and motility traits. Our collection of 962 metagenome-assembled genomes presents the largest collection of pelagic ODZ microbes and reveals a clearer picture of the nitrogen cycling community within this environment.

## Introduction

Bioavailable (i.e., fixed) nitrogen limits biological productivity in much of the surface ocean, thus the processes that balance the nitrogen budget require a thorough understanding [1]. Microbes mediate the biogeochemical transformations of nitrogen in its various forms, including the removal of fixed nitrogen via denitrification. A process of anaerobic respiration in which nitrogen oxides act as terminal electron acceptors, denitrification occurs when oxygen is limiting for aerobes. Three major oceanic oxygen deficient zones (ODZs), located in the eastern tropical North Pacific (ETNP), the eastern tropical South Pacific (ETSP), and the Arabian Sea, account for nearly all of water column fixed nitrogen loss through the processes of denitrification and anaerobic ammonium oxidation (anammox) and about 30% of total oceanic fixed nitrogen loss despite containing only 0.1–0.2% of oceanic volume [2]. Hotspots of denitrification, ODZs harbor diverse microbial assemblages for which much remains to be understood regarding their genetics, despite their disproportionate contributions to marine nitrogen cycling [3–5].

Many previous studies on denitrifying communities within ODZs have largely relied upon PCR amplification of denitrification marker genes [3, 6–9]. A stepwise process, denitrification is driven by the action of enzymes encoded by a suite of known genes (Figure 1). The first step of nitrate (NO_3_^−^) reduction to nitrite (NO_2_^−^) occurs via the periplasmic NO_3_^−^ reductase encoded by *nap* or the membrane-bound NO_3_ reductase encoded by *nar*. The second step, NO_2_^−^ reduction to nitric oxide (NO), is catalyzed by two functionally similar but structurally distinct NO_2_^−^ reductases, a copper-containing NO_2_^−^ reductase and a cytochrome *cd_1_-*containing NO_2_^−^ reductase encoded by *nirK* and *nirS*, respectively. The reduction of NO to nitrous oxide (N_2_O) proceeds through NO reductases encoded by diverse *nor* genes [10–12], and N_2_O is finally reduced to inert N_2_ gas by N_2_O reductases encoded by *nos* [13]. Despite the breadth of knowledge on denitrification in cultured isolates, recent sequencing advances have revealed wide diversity in denitrification genes missed by current PCR primers [12, 14–16]. Additionally, denitrification has been demonstrated to be a modular process, with some microbes able to perform the complete reduction of NO_3_^−^ to N_2_ while others are only capable of one or a subset of steps [17]. Increasingly, sequencing efforts demonstrate that diverse partial denitrifiers comprise the majority of environmental denitrifiers, yet denitrification is often treated as a complete, singular process in marine biogeochemistry [6, 18–21].

**Figure 1.**
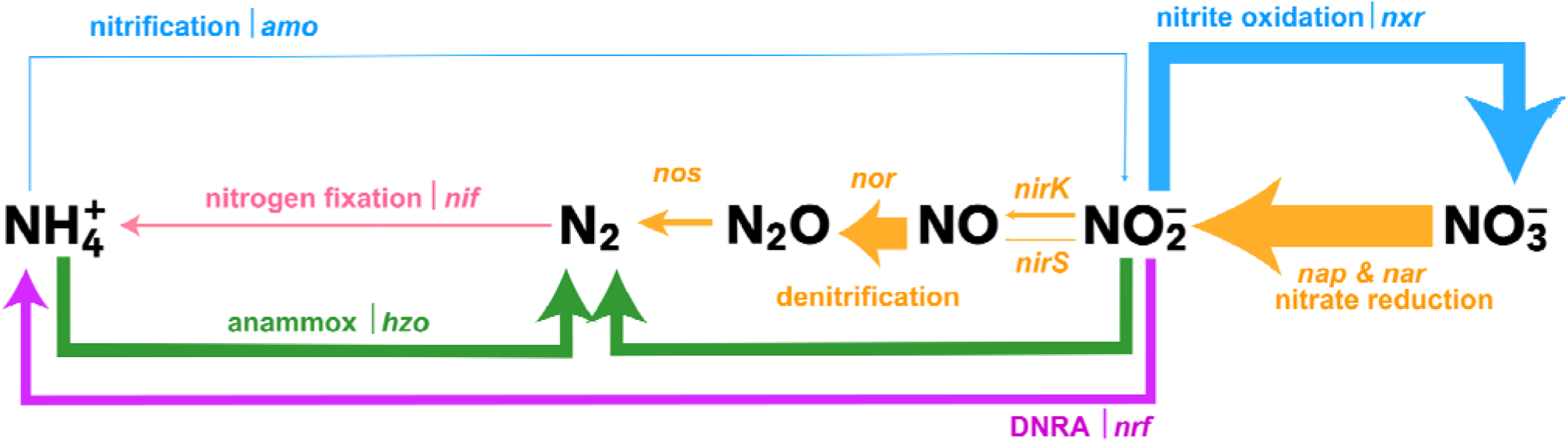
A schematic of nitrogen cycling within ODZ depths (O_2_ < 5 μM). Thickness of arrows correlates to total relative abundance of MAGs with the gene encoding the enzyme for that pathway, averaged across ODZ depths in the ETSP, ETNP, and Arabian Sea. Genes queried as proxies for each metabolism are italicized next to the name of the metabolism. Figure modified from Babbin et al. 2021 [98].

Partial denitrifiers hold important implications for biogeochemical cycling, as they may either produce or consume key denitrification intermediates depending on their specific genetic makeup. Denitrification intermediates interface with various other nitrogen metabolisms; for example, NO_2_^−^ participates in pathways for dissimilatory nitrate reduction to ammonia (DNRA), NO_2_^−^ oxidation to NO_3_^−^, and anammox (Figure 1). Furthermore, the emission of N_2_O, a major greenhouse gas, has been correlated with the presence of partial denitrifiers lacking *nos*, while N_2_O consumption might correlate with the presence of *nos*-carrying partial denitrifiers [14, 22]. ODZs are a major source of N_2_O, yet the magnitude and net effect of potential N_2_O emission and consumption within ODZs remains debated [7, 23–25].

Some previous estimates of the marine nitrogen budget indicate the prevalence of denitrification and anammox results in an excess of fixed nitrogen losses over gains [2, 26, 27], although these estimates include large uncertainties. Moving towards a better understanding of the marine nitrogen budget requires constraining these uncertainties by recognizing the relative contributions of nitrogen loss processes, nitrogen fixation, and fixed nitrogen recycling via nitrification, NO_2_^−^ oxidation, and DNRA, as well as which organisms perform these processes and the spatial coupling among them. However, difficulties culturing environmental microbes and incomplete primer coverage have limited our knowledge of the distribution, taxonomy, and abundance of ODZ denitrifiers [16]. We use genome-resolved metagenomics to assemble the largest collection of uncultured microbial genomes from global ODZs to date. Assessing these metagenome-assembled genomes (MAGs) for denitrification and other nitrogen cycling genes reveals the taxonomic identity of ODZ denitrifiers, their nitrogen cycling capabilities within the water column, and their potential impacts on biogeochemistry.

## Materials and Methods

### Sampling, Sequencing, and Read Quality

Sampling and sequencing methods for public ETNP metagenomes are described in Fuchsman et al. (2017), Glass et al. (2015), and Tsementzi et al. (2016). Sampling and sequencing methods for public ETSP metagenomes are described in Stewart et al. (2012) and Ganesh et al. (2014). Raw samples were downloaded from the Sequence Read Archive (SRA) using NCBI BioProject ID numbers: PRJNA350692 (Fuchsman et al. ETNP metagenomes), PRJNA254808 (Glass et al. ETNP metagenomes), PRJNA323946 (Tsementzi et al. ETNP metagenomes), PRJNA68419 (Stewart et al. ETSP metagenomes), and PRJNA217777 (Ganesh et al. ETSP metagenomes) (Table S1). Raw reads were trimmed with Trimmomatic v0.39 to remove adapters and low-quality bases using the LEADING:3 TRAILING:3 SLIDINGWINDOW:4:15 MINLEN:36 flags [32].

For 2016 ETNP, 2018 ETNP, and 2007 Arabian Sea metagenomes reported here for the first time, reads were sequenced at the DOE Joint Genome Institute (JGI) on an Illumina NovaSeq using paired end reads 151 base pairs in length. BBDuk v38.45 was used to remove contaminants, trim adapters, and remove low quality bases. Reads mapping to human and animal references as well as common microbial contaminants were removed using the JGI pipeline, and final reads were retrieved from JGI.

Sampling locations for each metagenome were visualized using Python 3.7.12 and the cartopy package. These were plotted against global oxygen concentrations from 300 m below sea surface from Ocean Data Atlas 2018 (Figure 2a). Data for vertical profiles of O_2_, NO_3_^−^, and NO_2_^−^ were retrieved from the original studies when available or from BCO-DMO (Figure 2b, S1). From NO_3_^−^, NO_2_^−^, and PO_4_^3−^ data, N* was calculated using the equation N*= (NO_3_^−^ + NO_2_^−^ – 16 *×* PO_4_^3−^) + 2.9 μmol kg^−1^ [33]. Oxygen measurements were made using a rosette containing a Conductivity-Temperature-Depth profiler equipped with a Seabird Clark-type dissolved oxygen electrode with a resolution in the micromolar oxygen range. While previous measurements with high resolution trace oxygen sensors show oxygen below the detection limit of 10 nM in ODZ cores [34, 35], the gradients exhibited by Clark-type electrodes remain valid for identifying ODZ waters [36]. Due to this resolution, we define ODZ depths as those with oxygen < 5 μM. Vertical profiles for O_2_, NO_3_^−^, NO_2_^−^, and N* were graphed using R ggplot2. Due to sparsity of NO_3_^−^ and NO_2_^−^ data for the ETNP and ETSP, profiles were aggregated to create a composite profile for each study, and smoothed using geom_smooth with the LOESS method in R ggplot2.

**Figure 2.**
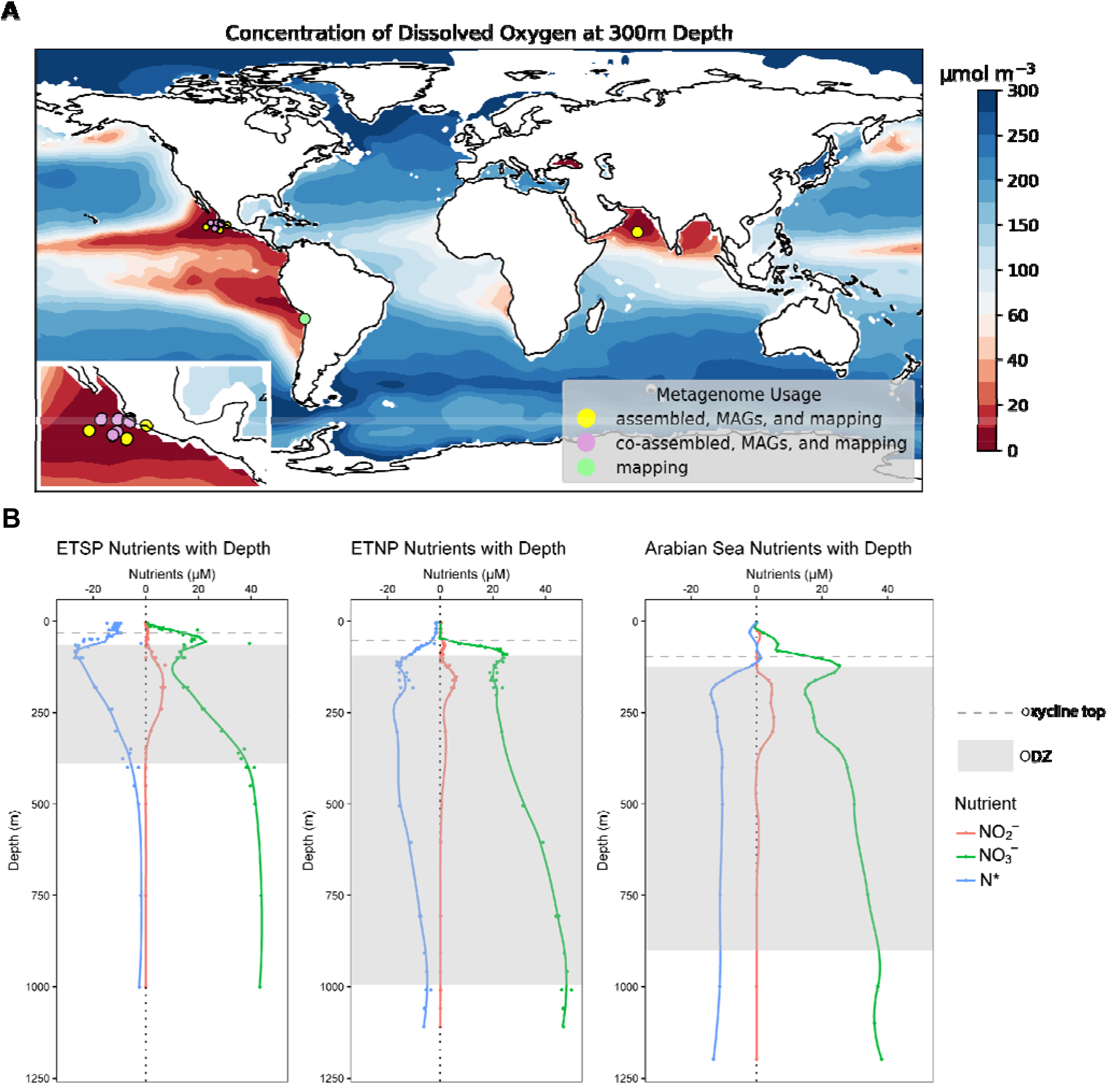
a) Locations of metagenomes from ETNP, ETSP, and Arabian Sea b) Representative profiles for NO_3_^−^, NO_2_^−^, and N* for all 3 major ODZs. ETSP and ETNP profiles are composite profiles from all sampling sites in Ganesh et al., 2014 and Fuchsman et al., 2017, respectively, while Arabian Sea profile is from sampling station 2 in Bulow et al., 2010, corresponding to Arabian Sea metagenomes used in this study. Dotted grey horizontal line indicates the top of the oxycline at each station based upon corresponding O_2_ CTD data, and shaded grey box indicates the ODZ (O_2_ < 5 μM). O_2_ profiles are from Ganesh et al., 2014 sampling station 1, ETNP profile is from Fuchsman et al., 2017 sampling station 141, Additional O_2_ profiles can be found in Supplementary Figure 1.

### Metagenome Assembly and Binning

Reads were assembled with MEGAHIT v1.2.9 [37] using default parameters. For metagenomes from the 2016 ETNP, 2018 ETNP, and 2007 Arabian Sea collections, each metagenome was assembled separately. For the Fuchsman et al. ETNP 2012, Glass et al. ETNP 2013, and Tsementzi et al. ETNP 2013 metagenomes, all samples from the same study were co-assembled to maximize read depth. Co-assembly was performed for ETSP metagenomes, but low read depths prevented successful binning and further analysis. Short reads from ETSP metagenomes were retained for mapping to our final set of MAGs.

From each ETNP and Arabian Sea assembly, contigs shorter than 800 nucleotides were removed, Bowtie v2.3.5 was used to map reads to assemblies, and samtools v1.7 sorted and indexed the resultant bam files. Binning was performed using CONCOCT v1.00 [38], Metabat2 v2.12.1 [39], and Maxbin2 v2.2.6 [40] within the metaWRAP v1.3 wrapper [41]. From the three bin sets per assembly, the best non-redundant bins were selected using the metaWRAP bin_refinement module. Resulting bins were further improved with the metaWRAP reassemble_bins module, in which reads belonging to each bin are extracted and reassembled with both a “strict” and “permissive” algorithm, and only bins which are improved by this reassembly are altered in the final set of bins. Bin quality was assessed using CheckM v1.0.12 and medium to high quality metagenome-assembled genomes (MAGs) were defined as bins with completion >50% and contamination <10% [42].

### Gene searching within Metagenomes

For each metagenome, assemblies were annotated with PROKKA v1.14.6 against the HAMAP [43] and Pfam databases [44]. Annotated assemblies were queried with custom HMMs using HMMer3 [45] for the denitrification genes *napA, narG, nirK, nirS*, and *nosZ*, as well as the DNRA gene *nrfA*, the nitrogen fixation gene *nifH*, the NO_2_^−^ oxidation gene *nxrA*, the nitrification gene *amoA* for ammonium-oxidizing archaea and bacteria separately, and the anammox gene *hzo*. For NO reductases, individual HMMs for canonical *qnor* and *cnor* variants, along with HMMs for non-canonical *bnor, enor, gnor, nnor*, and *snor* genes and the NO dismutase gene *nod*, were downloaded from Murali et al., 2021 [10] and used as queries. Furthermore, assemblies were searched using custom HMMs for the single-copy genes (SCGs) *rpoB, rplB, gyrB, rpS3*, and *recA*, which are conserved in bacteria and archaea. Normalization was performed by dividing the numbers of hits for each nitrogen cycling gene by the average number of hits for the five SCGs per metagenome assembly. Pearson correlations were done for all normalized percentages of nitrogen cycling genes with each other and with NO_3_^−^, NO_2_^−^, and depth. The corr.test function in the R psych package v2.2.9 was used to calculate the Pearson correlations and test statistics (p-value) using a two-tailed t-test to determine if correlations differ significantly from 0. We corrected for multiple hypothesis testing with the Benjamini-Hochberg method [46], and a significance threshold of p < 0.01 was chosen. Correlation heatmaps were visualized with the R heatmaply package 1.4.0.

### MAG Analyses, Mapping, and Gene Searching

Taxonomy was assigned to all MAGs using GTDB-tk v1.7.0 [47] with the classify_wf workflow. MAGs were annotated with PROKKA v1.14.6 [48] against the HAMAP [43] and Pfam databases [44]. For coverage mapping and comparison purposes, MAGs were dereplicated with dRep v3.2.2 with the -sa 0.99 flag to produce a final collection of non-redundant genomes. Coverage mapping was performed with CoverM on dereplicated genomes using the flags minimap2-sr --min-read-aligned-percent 50 --min-read-percent-identity 0.95 --min-covered-fraction 0 (https://github.com/wwood/CoverM). Relative abundances of dereplicated MAGs resulting from CoverM mapping were visualized using R v4.1.3 and the packages phyloseq, ggplot2, and dplyr (Figure S2).

We minimized the risk of protein misidentification (false positives) while maximizing the retrieval of non-canonical genes or more distant homologues using curated HMMs along with examination of gene hits for the presence of conserved active site residues. PROKKA annotations for each MAG were queried for nitrogen cycling genes using custom HMMs as described above. From positive hits for each gene, the corresponding protein sequences were retrieved and multiple sequence alignments generated with MAFFT v7.450 using the --auto and --leavegappyregion parameters. These alignments were visualized in JalView v2.11.2.6 [49] and inspected for alignment quality and the conservation of active site amino acid residues for each protein [9, 10, 16, 50–55]. Validated hits were used to generate diagrams of gene presence and absence for each MAG. The number of MAGs belonging to each nitrogen cycle gene combination was visualized using R v4.1.3 and the UpSetR package. Additionally, relative abundances of MAGs carrying each gene across all metagenomes were visualized with the R packages phyloseq, ggplot2, and dplyr. Euler diagrams showing relative abundances of MAGs carrying each denitrification step for each metagenome were calculated by adding up relative abundances of all MAGs capable of each of the four denitrification steps, and visualized with the R eulerr package.

For bacterial motility, chemotaxis, and aerotaxis genes, custom HMMs were downloaded from Hallstrøm et al., 2022. The chemotaxis genes *cheA, cheB*, and *cheR*, the aerotaxis receptor gene *aer*, and the genes *fliG, fliM*, and *fliNY* involved in bacterial flagellar motion were searched against all MAGs. Chemotaxis or motility were considered present if at least 2 out of the 3 queried chemotaxis or motility genes were present in the MAG, to account for varying MAG completeness. Additionally, sequences for the archaeal flagellar gene *flaJ* were downloaded from the NCBI conserved domain database (CDD) and aligned with MAFFT v7.450 using the mafft-linsi and --leavegappyregion. HMMs were built for *flaJ* and searched against all MAGs, and archaea possessing this gene were considered motile.

## Results

### Biogeochemical context

The distribution of denitrifying genes and MAGs was investigated in 56 metagenomes spanning multiple depths, years, sites, and studies from the three permanent, open ocean ODZs (Figure 2a). Oxygen profiles corresponding to each metagenome show a characteristic rapid decrease from surface saturation to anoxia, termed the oxycline, occurring approximately between 50– 100m depth (Figure 2b, S1). Below the oxycline, oxygen levels remain beneath the detection limit until around 500m below the sea surface for the ETSP and 1000m below the sea surface for the ETNP and Arabian Sea. Corresponding profiles of NO_3_^−^ and NO_2_^−^ reveal typical decreases in NO_3_^−^and increases in NO_2_^−^ (the secondary nitrite maximum, or SNM) in this region (Figure 2b). N* values become increasingly negative within the oxycline and ODZ, indicating a net nitrogen deficit relative to phosphorus characteristic of the nitrogen loss processes denitrification and anammox.

Within the mixed, oxygenated layers above the ODZ, denitrification genes in the individually assembled metagenomes are scarce, but increase in abundance at the deeper hypoxic and anoxic waters of the oxycline and ODZ (Figure S3a). While the *nirK* gene, encoding copper-containing NO_2_^−^ reductase, peaks in abundance at 100 m below sea surface, the *nirS* cytochrome *cd*_1_-containing NO_2_^−^ reductase, *nor* NO reductase, and *nosZ* N_2_O reductase genes peak at approximately the same depths, below 150m. Pearson correlations reveal a significant positive correlation among these three genes (p<0.01), along with the NO_3_^−^reductase genes *napA* and *narG*, the anammox hydrazine oxidoreductase gene *hzo*, the NO_2_^−^ oxidation gene *nxrA*, and the DNRA gene *nrfA*. In contrast, *nirK* exhibits little correlation with other denitrification genes, yet correlates positively with the *amoA* gene for ammonia monooxygenase (p=0.003) and negatively with NO_2_^−^ (p=0.004) and oxygen (p=0.003) (Figure 3a).

**Figure 3.**
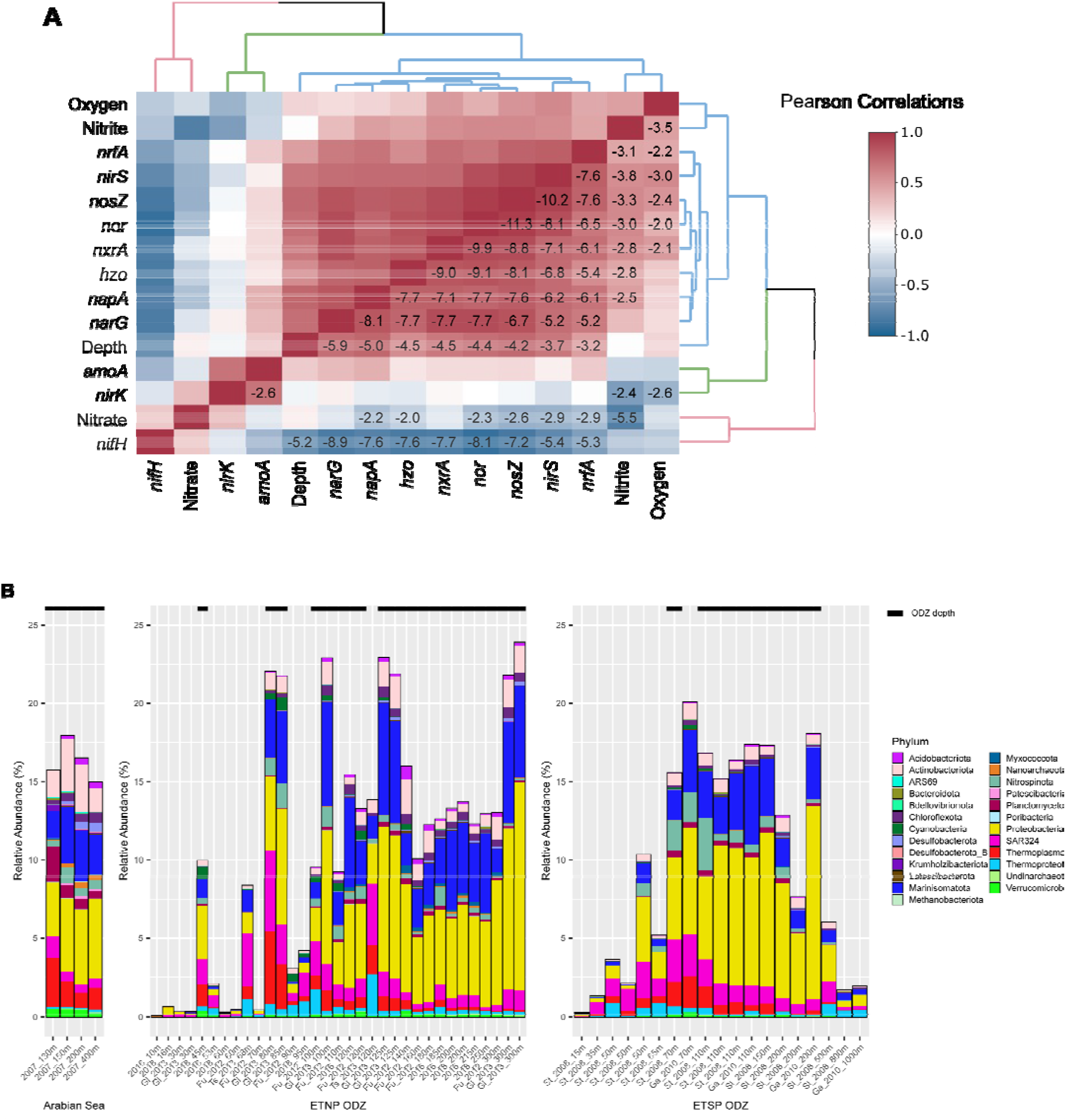
a) Pearson correlations for all relevant nitrogen cycling genes with p-values overlaid, showing only log_10_-transformed significant p-values (p < 0.01) after a Benjamini-Hochberg multiple hypothesis correction. b) Relative abundances of denitrifier MAGs across metagenomes, color-coded by phylum-level taxonomy, from the three major ODZs. Black bars above the graph indicate ODZ depths (O_2_ < 5 μM).

### ODZ MAG collection and denitrifying community stability

To determine if these positive correlations between *napA, narG, nirS, nor*, and *nosZ* are due to co-occurring organisms or co-occurrence of these genes within the same organisms, we generated 962 MAGs >50% completion and <10% contamination from the ETNP and Arabian Sea, with 245 high-quality MAGs >90% completion. After dereplication, these MAGs contain 307 uncharacterized taxa at the species level out of 497 non-redundant genomes, including 1 MAG novel at the phylum level, 2 at the class level, 8 at the order level, and 19 at the family level (Supplementary Table S1), as determined by the lowest level taxonomic assignment performed by GTDB-tk. Mapping non-redundant MAGs from our collection to all three global ODZs reveals a distinctive and diverse ODZ community that is stable at the phylum level across time, sampling sites, and ODZs (Figure S2). Within ODZ depths, up to approximately 50% of assembled reads map to MAGs, while mapping rates are lower above and below the ODZ. In anoxic layers, Proteobacteria are the most abundant, followed by Marinisomatota. Phyla present in lower abundance across most or all metagenomes include Actinobacteria, Planctomycetes, SAR324, Nitrospinota, and Thermoplasmatota (Figure S2).

Closer examination of the denitrifying community, defined as MAGs harboring at least one denitrification gene, reveals a similar pattern of phylum-level abundance, distribution, and stability across the ETSP and ETNP (Figure 3b). Arabian Sea metagenomes also reveal similar proportions as the ETNP and ETSP at similar depths, but the limited number of sampling depths restricts comparison. Denitrifier draft genomes represent up to 24% of the ODZ microbial community by relative abundance, and likely more as 50% or more of the community did not map to our MAG collection. Proteobacteria and Marinisomatota dominate within denitrifiers, but the denitrifying community includes 22 of the 34 total phyla into which all MAGs were assigned (Figure 3b).

### Denitrifying phyla and relative abundances within the ODZ water column

Single-step *napA* or *narG* NO_3_^−^ reducers, single-step *nor* NO reducers, and single-step *nosZ* N_2_O reducers dominate in number of MAGs. 105 ODZ MAGs (11%) have *narG*, with 59 (58% of narG MAGs) possessing no other denitrification genes (Figure 4a). Similarly, *napA* MAGs account for 109 MAGs total, and 60 (55%) of these have only *napA*. The overlap of *napA* and *narG* in MAGs is rare (14 MAGs). A total of 200 (21%) MAGs have NO_3_^−^ reduction capability, and the majority of both types of NO_3_^−^ reducers possess no further denitrification capabilities (Figure 4). In terms of relative abundance, *napA* NO_3_^−^ reducers reach about 5% of the community in the ODZ core, while *narG* NO_3_^−^ reducers reach about 12%, (Figure 5a, S5a). While *narG* nitrate reducers coexist with *napA* NO_3_^−^ reducers across depths, Proteobacteria and SAR324 dominate within *napA* NO_3_ reducers, while Marinisomatota and Proteobacteria comprise the most abundant fractions of *narG* NO_3_^−^ reducers. Within metagenome assemblies, *narG* comprises up to 40% of gene hits normalized to a set of single-copy housekeeping genes (SCGs) (Figure S3a), the most abundant gene within our queried set. *NapA* is the second most commonly found denitrification gene within the metagenomes at around 30% abundance compared to our SCGs, while *nirS* and *nirK* comprise 10% or less at ODZ depths (Figure S3a).

**Figure 4.**
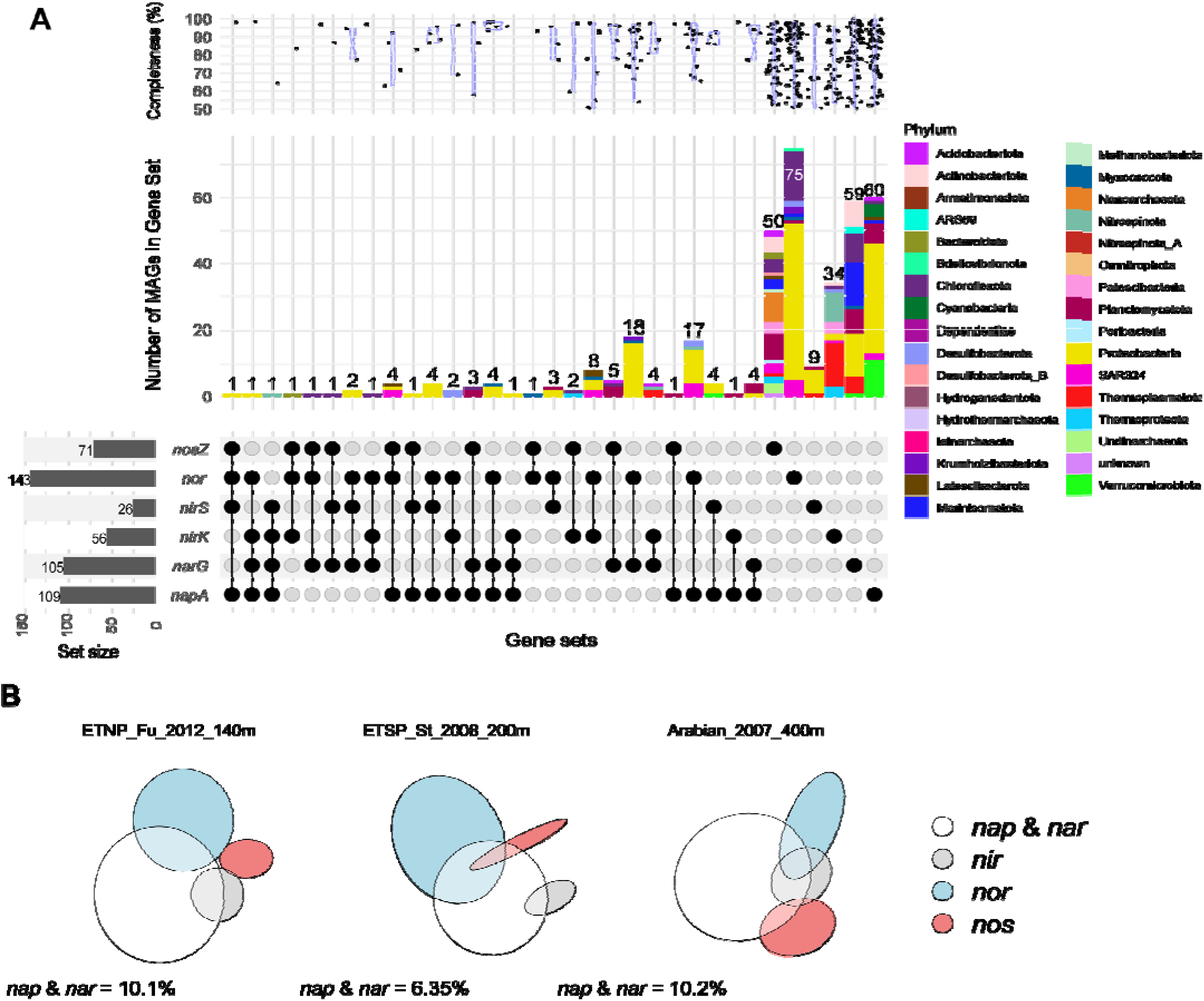
a) The number of MAGs carrying each denitrification gene set within the ODZ MAG collection. Top panel shows genome completion distributions for MAGs belonging to each gene set, middle panel shows the number of MAGs colored by phylum-level taxonomy, and bottom panel shows the genes within each gene set. Left bottom panel shows the number of MAGs carrying each specific nitrogen cycle gene. Additional graphs showing all nitrogen cycling genes can be found in Supplementary Figure S4. b) Representative Euler diagrams for ODZ depths showing the relative abundance of the four steps of denitrification. Circles and intersections are scaled to the total relative abundance of all MAGs possessing the genes for that step or step combination. The white circle corresponds to the relative abundance of MAGs with *napA, narG*, or both within that metagenome. All Euler diagrams for all metagenomes can be found in Supplementary Figure S8.

**Figure 5.**
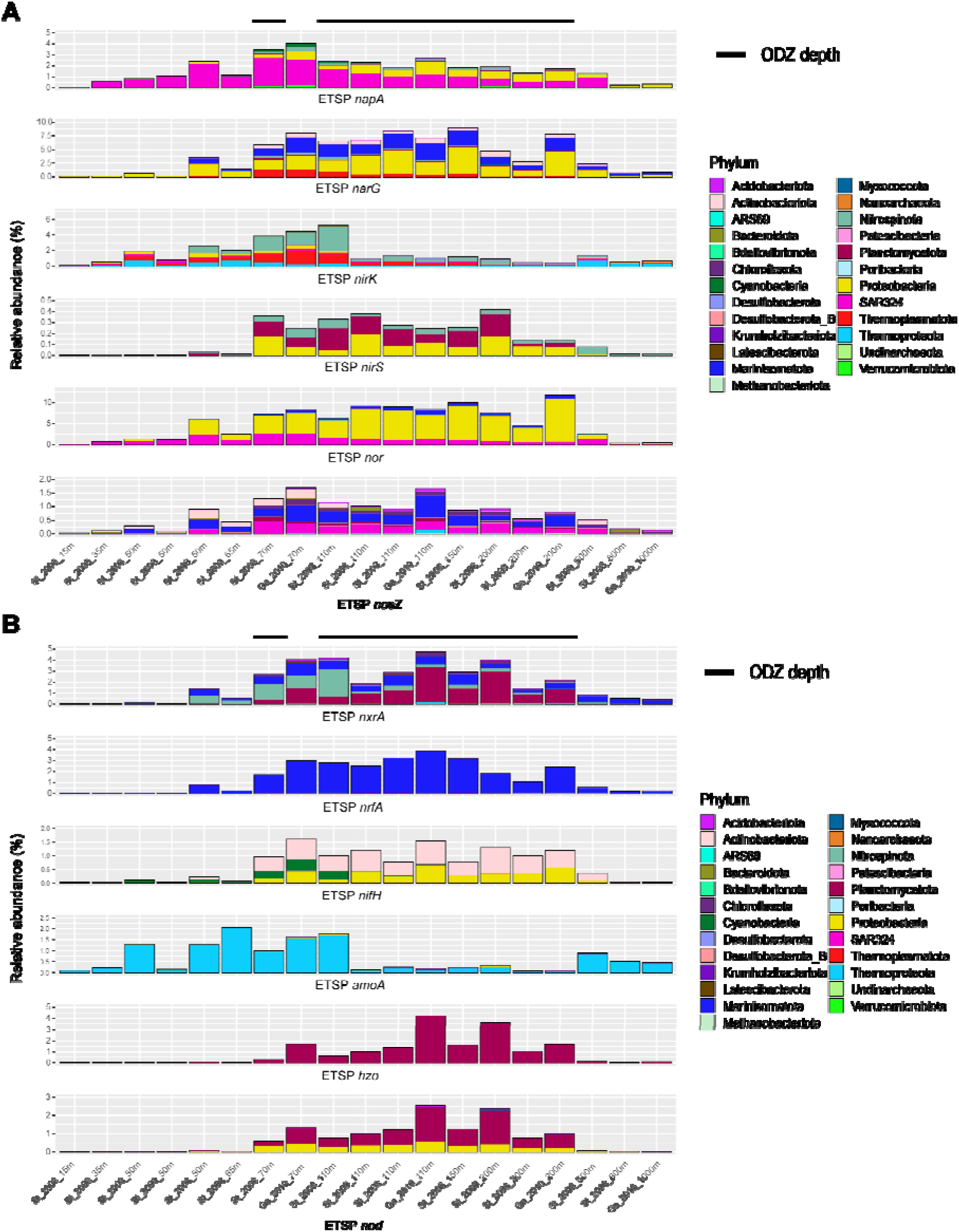
a) Relative abundance mapping of MAGs carrying each denitrification gene across all ETSP metagenomes. MAGs are colored by phylum-level taxonomy. b) Relative abundance mapping of MAGs carrying each non-denitrification nitrogen cycling gene across all ETSP metagenomes. MAGs are colored by phylum-level taxonomy. Black bars above the graph indicate ODZ depths (O_2_ < 5 μM). Additional graphs for ETNP metagenomes can be found in Supplementary Figure S5.

Consistent with the relatively high abundance of *nor* in metagenomes (up to 25% at ODZ depths), 143 MAGs carried a variety of *nor*, with 75 (53%) possessing no other denitrification genes (Figure 4a). However, of these only 62 carry the canonical variants *qnor* and *cnor*, while 91 carry a non-canonical *nor* (Figure S6). Total relative abundance of all *nor* MAGs reaches up to 15% at ODZ depths (Figure 4a, S5a), but MAGs carrying non-canonical *nor* reach higher relative abundance in both the ETNP and the ETSP compared to MAGs carrying *qnor* and *cnor* (Figure S6). Proteobacteria, SAR324, and Marinisomatota are the predominant NO reducers by relative abundance. For the 71 N_2_O reducers, 50 (70%) carry *nosZ* only (Figure 4a), and of the nosZ multi-gene denitrifiers (21 MAGs), the majority (17 MAGs) carry *nosZ* and *napA* or *narG*. *NosZ* MAGs account for up to 2% of the community while in metagenomes *nosZ* reaches about 15% of the community. Marinisomatota comprise a large fraction of N_2_O reducers by relative abundance, with smaller contributions by SAR324, Actinobacteria, and other phyla (Figure 4a, 5a, S5a).

NO_2_^−^ reducers carrying *nirK* account for 56 MAGs and up to 6% of the population by relative abundance at the top of the ODZ and in the oxycline, but drops to 2% within the deeper ODZ (Figure 5a, S5a). This pattern is also reflected within SCG-normalized gene counts in the metagenome assemblies, in which *nirK* reaches up to 40% of the SCG count at 100m but about 10% below 100m (Figure S3a). The most abundant *nirK* containing organisms within our MAGs belong to Nitrospinota, Thermoplasmatota, and Thermoproteota, while *nirS* nitrite reducers are largely Proteobacteria and Planctomycetota (Figure 5a, S5a). We find only 9 *nirS* single-step denitrifiers (35%), compared to 34 (61%) *nirK* (Figure 4a). Out of 26 total *nirS* MAGs, 11 also carry *napA*, 3 carry *narG*, and 10 carry *nor*. However, with the exception of *nirS* denitrifiers, few overlaps exist between denitrifiers carrying genes for NO_2_^−^ reduction, NO reduction, and N_2_O reduction, with most denitrifiers specializing in only one of these steps (Figure 4b, S8). Out of 383 denitrifying MAGs, we obtained only a single complete denitrifier belonging to Alphaproteobacteria.

### Other nitrogen cycling phyla, relative abundances, and distributions

Surprisingly, MAGs encoding *nxrA*, the nitrite oxidoreductase gene for oxidizing NO_2_^−^to NO_3_^−^, accounted for 111 MAGs, 43 of which carried only *nxrA* (Figure S4b). 38 MAGs also harbored *nxrA* and *nrfA*, accounting for 88% of *nrfA* MAGs total. Other nitrogen cycling genes examined, *nifH*, *hzo*, and *amoA*, did not tend to co-occur with denitrification genes in MAGs (Figure 3, S4b), although 2 *Candidatus* Scalindua MAGs carrying *hzo* also carried *nirS*, in line with previous reports of anammox *nirS* [56]. A previous amplicon-based survey indicated low *nirS* abundance and diversity in ODZs, with the most abundant *nirS* OTU similar to *nirS* from *Ca*.Scalindua [57]. These *Ca*. Scalindua MAGs lack a *nor* gene, but two *Ca*. Scalindua MAGs carry the *nod* gene for NO dismutase. Relative abundances reveal that *nxr* MAGs and *hzo* MAGs each comprise up to 8% of the ODZ community, *nrfA*-carrying MAGs up to 6%, and *nifH* and *amoA* MAGs up to around 3% each (Figure 5b, S5b). The single-copy-gene-normalized metagenome data largely supports these trends, although a higher percentage of *nxrA* genes (up to 40%) compared to *hzo* genes (up to 15%) were found within metagenomes (Figure S3b). While surface metagenomes show a near-zero count of denitrification and most other nitrogen-cycling genes, *nifH* counts peak at the surface and quickly drop to near zero below 100m depth. As few of our MAGs were assembled from or mapped to surface metagenomes (Figure S2), our MAG-based estimates of *nifH* underestimate the number and abundance of nitrogen fixers.

Despite the diversity of MAGs carrying the *nrfA* and *nxrA* genes, the majority of the *nxrA* community in the ETNP and ETSP are Planctomycetota, with smaller fractions contributed by Nitrospinota, Marinisomatota, and Chloroflexota. The *nrfA* community is almost entirely dominated by Marinisomatota by relative abundance. Similarly, *amoA* is dominated by Thermoproteota, ammonia-oxidizing archaea (AOA). Nitrogen fixers carrying *nifH* in the upper depths of the ODZ and the oxycline are primarily Cyanobacteria, while deeper depths are dominated by heterotrophic Proteobacteria and Actinobacteria. As expected, all *hzo* genes in our dataset belong to Planctomycetes assigned to *Ca*. Scalindua, the dominant ODZ anammox organism, which despite numbering only 5 MAGs in this set comprises up to 4% and 9% of relative abundance in the ETSP and ETNP ODZ communities, respectively (Figure 5b, S5b).

Overall relative abundances of nitrogen cycling genes, calculated using both SCG-normalized metagenome hits and MAG relative abundance mapping, follow similar patterns. The potential for NO_3_^−^ reduction to NO_2_^−^ dominates across almost all ODZ depths (Figures 4b, S3, S8). The next largest contribution is potential *nor*-mediated NO reduction to N_2_O. However, *nirK*-catalyzed NO_2_^−^ reduction potential peaks between 50-100m in the oxycline and upper ODZ, but forms a much smaller proportion below 100m (Figure S3a, S8). While contributions of *amoA* and *nifH* to the ODZ community are smaller, the genes and organisms are not absent (Figure 5b, S5b). MAGs harboring the *nrfA* gene are consistently present across ODZ depths. The potential for NO_2_^−^ oxidation to NO_3_^−^ mediated by *nxr* appears substantially in both metagenomes and MAGs, and is widespread across MAGs.

### Chemotaxis and motility genes in denitrifying MAGs

Searching for motility and chemotaxis-related genes within MAGs reveals a higher proportion of these traits within denitrifiers compared to non-denitrifiers (Figure 6a, Figure S7a). Within 383 denitrifying MAGs, 108 (28%) possessed chemotaxis, motility, or aerotaxis capability, compared to19% of non-denitrifiers. Comparing denitrifiers to non-denitrifiers, 20% vs. 12% were motile, 14% vs. 5% chemotactic, and 8% vs. 2% aerotactic. MAGs with both chemotaxis and motility genes were particularly prevalent in the denitrifying community, comprising 20–50% in the oxycline and upper ODZ (Figure 6a), while at the same depths they are less than 5% of the non-denitrifying community (Figure S7a). Less than 1% of non-denitrifiers have both aerotaxis and chemotaxis or aerotaxis and motility, but 5–20% of denitrifiers have these traits at ODZ depths, with an even higher percentage in the oxycline. However, motility without chemotaxis or aerotaxis appears widespread among non-denitrifiers (Figure S6a).

**Figure 6.**
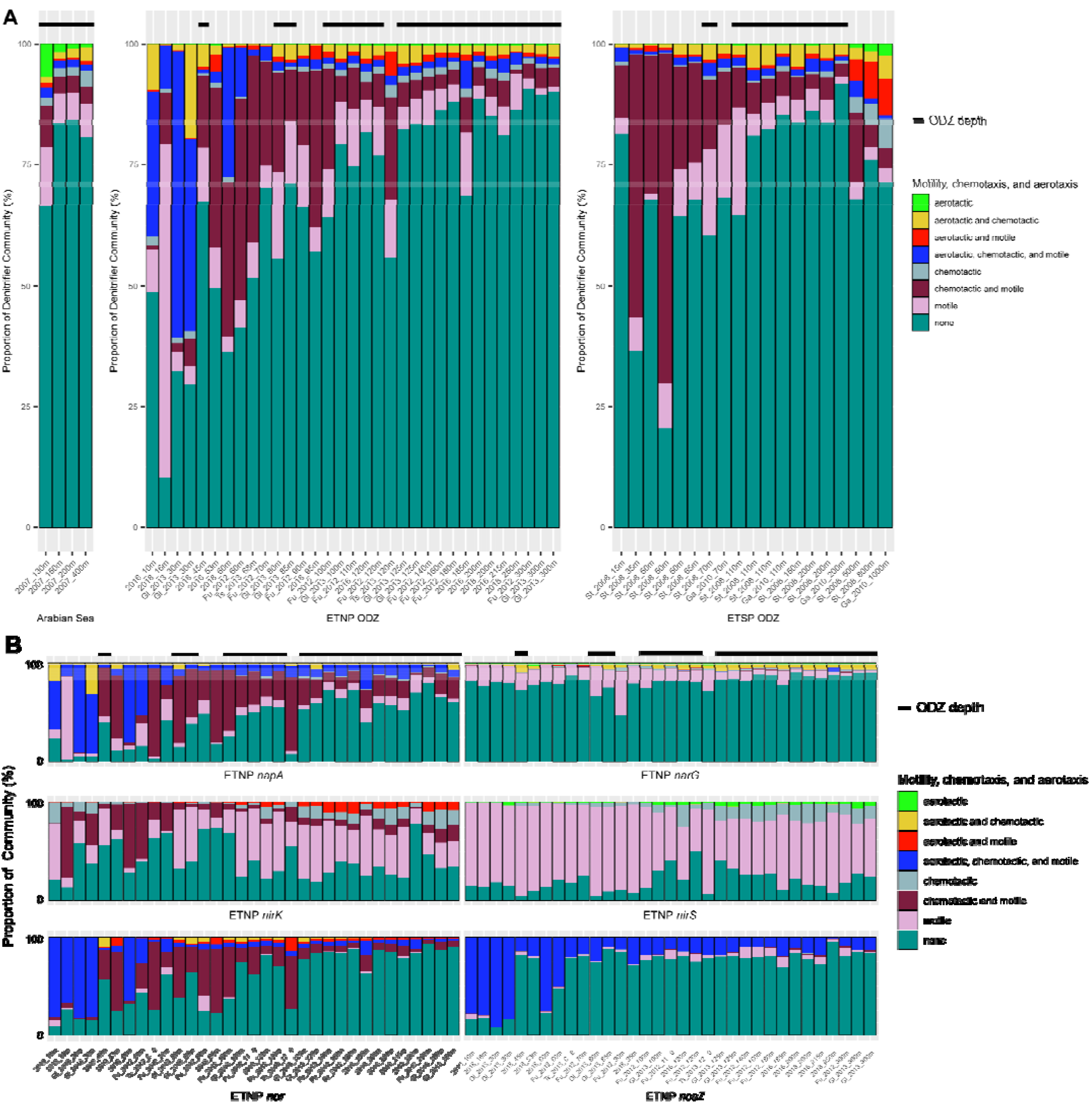
a) Relative abundances of denitrifier MAGs across all metagenomes, color-coded by presence of motility, chemotaxis, and aerotaxis genes. Relative abundances of non-denitrifying MAGs showing presence of motility, chemotaxis, and aerotaxis genes can be found in Supplementary Figure S6a. b) Relative abundance mapping of all MAGs carrying each denitrification gene across all ETNP metagenomes. MAGs are included if the denitrification gene in question is present, regardless of the presence or absence of other genes. MAGs are colored by the presence of motility, chemotaxis, and aerotaxis genes. Additional mapping results for ETSP metagenomes can be found in Supplementary Figure S6b.

Motile and chemotactic MAGs are in highest abundance within the *napA* community, making up the majority at oxycline and upper ODZ depths (Figure 6b, S7b). However, they make up very little (<1%) of the *narG* community, which is primarily dominated by non-motile organisms. Motile or chemotactic MAGs also comprise a sizeable fraction of *nirK, nirS, nor*, and *nosZ* communities, with *nirK* and *nirS* dominated by motile but not chemotactic or aerotactic MAGs, *nor* by motile MAGs with chemotaxis and/or aerotaxis, and *nosZ* by MAGs with aerotaxis, chemotaxis, and motility. With the exception of *nxrA* and *nod*, communities with the other nitrogen cycling genes are primarily non-motile, non-chemotactic, and non-aerotactic (Figure S7c).

## Discussion

### Unique ODZ microbial communities reflected by the largest MAG collection

Despite large-scale microbial sampling and sequencing efforts for the global oceans in recent years [58–62], fewer efforts have been conducted to date focusing on the open ocean oxygen-deficient zones. Yet, those efforts devoted to ODZ sequencing have revealed much about the microbial community therein, including marked shits in community structure from oxic surface waters to the anoxic core [30], diverse, atypical denitrification genes [12, 14], distinct particle-associated and planktonic assemblages [31], and a large number of uncultured organisms with surprising metabolic capabilities [63–66]. ODZs, due to their distinctive chemical profiles compared to the oxygenated ocean, harbor unique, largely uncultivated microbial communities [1]. Our 962 MAGs present the largest collection of genomes from global, open-ocean ODZs to date. The low mapping rate of our ODZ MAGs to surface metagenomes indicate a resident microbial community within the ODZ distinct from that of the surface ocean (Figure S2). This is consistent with previous studies using 16S amplicon sequencing, which report the clustering of ODZ and oxycline communities to the exclusion of surface communities based on community phylogenetic composition [31], as well as ODZ metagenomes not included in this study [63].

The increase in relative abundance of anaerobic nitrogen metabolism genes, such as those involved in denitrification and DNRA, from the surface to ODZ depths indicates adaptation of the microbial community to low-oxygen conditions. The paucity of ODZ genomes has historically hampered the taxonomic classification of ODZ nitrogen cycling microbes, as previous studies using reference genomes have reported a large number of nitrogen gene sequences unassigned to a particular bacterial or archaeal phylum [31]. By assembling genomes from metagenomes, we find that a stable denitrifying community dominated by several phyla. Proteobacteria, one of the most abundant phyla across ODZ depths, include a wide range of Alphaproteobacteria and Gammaproteobacteria known to participate in heterotrophic denitrification [31]. Another abundant ODZ phylum is the Marinisomatota, also known as Marine Group A, SAR406, or Marinimicrobia [64]. Despite being widespread in anoxic marine environments, little is known about the metabolism and ecophysiology of this uncultivated clade [64]. However, two Marinisomatota MAGs from anoxic waters of the Costa Rican Golfo Dulce were reported to contain and actively transcribe *nar* genes for NO_3_^−^ reduction [64], and a separate study of the ETSP found NO_3_^−^reductases in 6 out of 8 Marinisomatota MAGs, with 3 of these MAGs in the top 5 most abundant in the collection [69]. We confirm an important role for this clade as a NO_3_^−^ reducer within pelagic ODZs across all three major ODZs.

### Niche differentiation of denitrifiers carrying each gene

Despite co-occurrence among the denitrification genes *napA, narG, nirS, nor*, and *nosZ*, these genes are carried by different microbes rather than within the genomes of complete denitrifiers. Although we identified only a single complete denitrifier, this does not preclude the existence of complete denitrifiers in this environment since MAGs vary in completeness and may not capture the entire genome, and not all organisms assembled into MAGs. However, the presence of many partial denitrifiers with over 90% completeness (Figure S4a) and disparity between relative abundances of different denitrification genes in metagenomes (Figure S3a) point to the rarity of complete denitrifiers. Our results corroborate previous reports of partial denitrifier prevalence in agricultural and tundra soils [15, 70] as well as within a small number of previously-described ETSP ODZ MAGs [69].

While NO_3_^−^ reducers carrying *narG* and those carrying *napA* co-occur across depths, these genes do not frequently co-occur within genomes and are largely carried by different phyla. SAR324, formerly classified within the Deltaproteobacteria but recently reclassified into its own phylum, is the most abundant *napA* NO_3_^−^ reducer but largely absent in *narG* NO_3_^−^ reducers. A diverse clade with wide oceanic distributions, particularly in low-oxygen zones [71], little is known about this uncultivated phylum. SAR324 genomes from a hydrothermal plume carried *nosZ* and *nirK* [72], while other reported SAR324 genomes held minimal nitrogen cycling capabilities [73], pointing to the metabolic flexibility of this group. We find SAR324 to comprise 30% or more of *napA* NO_3_ reducers, *nor* NO reducers, and *nosZ* N_2_O reducers within ODZ depths by relative abundance. No *nap* or *nar* genes were discovered within 8 MAGs in our collection assigned to the SAR11 family Pelagibacteraceae, despite previous reports of SAR11 NO_3_^−^ reducers [29]. However, high genomic diversity and frequent recombination in SAR11 populations poses challenges for generating high-quality SAR11 MAGs [74], and only 3 of our SAR11 MAGs are >70% completion. Our MAG-based analysis likely underrepresents ODZ SAR11 abundance and diversity, and this may contribute to the higher relative abundance of *nap/nar* in metagenomes compared to the relative abundance of NO_3_^−^ reducer MAGs.

Although the enzymes encoded by *napA* and *narG* perform the same reduction process, the *napA* periplasmic nitrate reductase is not thought to provide energy for cells via a proton motive force, whereas the *narG* membrane-bound nitrate reductase does [75]. *NapA* has been implicated in balancing the intracellular redox state under oxygen limitation [76], so the metabolic context in which these two genes are used may also differ. However, *napA* does produce NO_2_^−^which can be used by *nir*, resulting in observable overlap between *napA* and *nirS*. Our results imply niche differentiation between these two types of NO_3_^−^ reducers. High NO_3_^−^concentrations in ODZs may enable co-existence of these NO_3_^−^ reducer ecotypes, particularly since NO_3_^−^is never depleted in the bulk ocean. Furthermore, abundant *napA* NO_3_^−^ reducers frequently carry motility and chemotaxis genes, which may facilitate particle colonization [77–79] and suggests a particle-associated lifestyle, whereas dominant *narG* NO_3_^−^ reducers are primarily non-motile (Figure 6b, S7b). The lack of motility among *narG*-containing organisms suggests a more planktonic lifestyle whereby these organisms are sustained by dissolved organic and inorganic nutrients. Moreover, both Ganesh et al. and Fuchsman et al. [12, 31] reported that *narG* is more frequently contained among planktonic cells rather than particle-associated size classes.

Similarly, niche differentiation appears between *nirS* and *nirK* nitrite reducers (Figure 5a, S5a). MAGs carrying *nirK* genes belong largely to ammonia oxidizing archaea (AOA) or nitrite oxidizing bacteria (NOB) involved in the nitrification pathway, including Nitrospinota, the dominant marine NO_2_^−^ oxidizer [80]. We find that *nirK* may predominate within nitrifiers and enable the process of nitrifier-denitrification [81]. The higher abundance of *nirK* vs. *nirS* points to nitrifier-denitrification as a potential source of N_2_O, particularly above 100m where a peak in *nirK* abundance concurs with a peak of N_2_O accumulation [82]. In contrast, *nirS* may predominate within heterotrophs and are more commonly multi-step denitrifiers. While *nirS* and *nirK* have been used as functional markers for denitrification [3, 83], these genes cannot be viewed interchangeably and may not reflect the activity of upstream or downstream denitrification steps.

Interestingly, the majority of *nir* MAGs, particularly those with *nirK*, lack *nor* (Figure 4), despite the toxicity of the resultant NO. Diverse, non-canonical *nor* genes have recently been discovered [10, 11, 84] and comprise the majority of *nor* genes we identified. Few *nir*-possessing MAGs also carried the NO dismutase gene *nod*, indicating novel NO detoxification methods likely exist within environmental microbes beyond canonical *nor* and *nod*. *NirK* in nitrifiers has been suggested to catalyze the oxidation of NO to NO_2_^−^ [85]. Alternatively, NO may act as an intercellular signaling molecule to modulate the behavior of interaction partners [86, 87]. The prevalence of *nor* genes, particularly single-step *nor* MAGs, may indicate a widespread need to detoxify NO despite the low concentrations of NO in bulk seawater, potentially as a result of NO secretion from spatially proximate neighbors such as within a particle. The diversity and function of non-canonical *nor* genes within ODZs has not previously been reported and warrants further exploration.

The prevalence of partial denitrifiers carrying only *nosZ* indicates potential for ODZ microbes to act as a sink for N_2_O, and supports previous work showing substantial consumption of N_2_O in oxyclines and ODZs [88] and rapid turnover of N_2_O [23]. However, the nanomolar concentrations of N_2_O, its rapid diffusion, and the low oxygen tolerance of N_2_O reductase poses challenges for denitrifiers relying solely upon *nosZ*. We find the motile fraction of *nosZ* MAGs to be dominated by MAGs with genes for aerotaxis, chemotaxis, and motility, comprising up to 30% of the *nosZ* community at ODZ depths (Figure 6, S7). The *aer*-encoded aerotaxis receptor senses oxygen gradients, and has been reported to facilitate negative aerotaxis, or the movement of cells away from oxygen [89, 90]. Along with chemotaxis, this may enable organisms to seek out localized regions of high carbon and N_2_O and low oxygen, such as particles. Previous metagenomic analyses of size-fractionated ETNP communities [12] found higher abundances of genes for the last two steps of denitrification on particles compared to free-living communities. Denitrification in particles is an active area of research, and has been posited to expand the niche of anaerobic metabolisms [91]. The higher abundance of motile and chemotactic denitrifiers, particularly ones carrying *nir, nor*, and *nosZ*, compared to non-denitrifiers (Figure 6, S7), supports the importance of particle-based denitrification and opens up fruitful avenues for further research into particle colonization and metabolisms.

The presence of 29 bacterial MAGs with aerotaxis or chemotaxis but not flagellar motility may result from MAG incompleteness, but the average completeness of 83% for these MAGs suggests the possibility of alternative motility mechanisms or functions of the chemosensory system. Potential interplay of the chemotaxis machinery has been described with pili-mediated surface motility, cell aggregation, virulence, and biofilm formation [92]. Further characterization of chemotaxis, aerotaxis, and motility in marine microbes presents an exciting route for future work.

### Other nitrogen cycling genes and the NO_3_^−^ ⇆ NO_2_^−^ loop

We find NO_2_^−^ oxidation potential in a diversity of phyla, yet only MAGs belonging to phyla Marinisomatota, Nitrospinota, and Planctomycetes reach relative abundances over 1% of the community (Figure 5b, S5b). NO_2_^−^ oxidation has been described as an aerobic process, but Nitrospinota have been discovered in ODZ waters and postulated to evolve from microaerobic or anaerobic ancestors [63, 80]. While the prevalence of NO_2_^−^ oxidation and discovery of Nitrospinota in ODZs has led to hypotheses of fully anaerobic Nitrospinota that use terminal electron acceptors other than oxygen for NO_2_^−^ oxidation, we find Nitrospinota dominant within *nxr*-carrying MAGs only at oxycline and upper ODZ depths (Figure 5b, S5b). Within permanently-anoxic ODZ depths, *nxr*-carrying Planctomycetes comprise the most abundant NO_2_^−^oxidizer, including MAGs belonging to the anammox bacterium *Ca*. Scalindua. The occurrence of *nxr* in anammox bacteria is well established [93, 94], and thought to harvest electrons for the reduction of NO_2_^−^to NO, an important intermediate in the anammox pathway. As high rates of NO_2_^−^ oxidation have been found in both oxycline and anoxic ODZ waters [95], microaerobic NO_2_^−^ oxidation by Nitrospinota and anaerobic NO_2_^−^ oxidation by anammox bacteria may both contribute to the NO_3_^−^ ⇆ NO_2_^−^ loop within ODZs.

Additionally, *nxr* co-occurs within MAGs along with various denitrification genes and the DNRA gene *nrfA* (Figure S4b) Possibly, *nxr* may confer a benefit to a wide variety of microbes by enabling autotrophic survival, but the low energy yield of aerobic NO_2_^−^ oxidation prevents this from being an efficient metabolism for growth. 38 out of 43 MAGs with *nrfA* also carry *nxr*, ranging across a diversity of phyla. The frequency of this co-occurrence suggests a benefit to organisms that can partition NO_2_^−^ to different metabolisms. Obtaining other nitrogen cycling genes may provide these microbes the metabolic flexibility to switch to higher-energy metabolisms when the situation allows. Previous work in the ETSP and Arabian Sea has uncovered substantial DNRA [56, 96], although they were unable to taxonomically link *nrfA* marker genes to taxonomic identity. We find *nrfA* widely distributed across phyla, but only Marinisomatota as an abundant DNRA organism in all ODZs and depths in which *nrfA* was present. The contribution of DNRA in ODZs and its interplay with nitrogen metabolisms such as NO_2_^−^ oxidation requires further exploration.

Previous studies have discussed the fate of NO_2_^−^ and whether it is primarily oxidized back to NO_3_^−^ or further reduced via downstream denitrification, anammox, or DNRA. Recent work using isotope measurements [6, 97] and proteomics [82] indicate a large contribution of NO_2_^−^ oxidation within anoxic waters. Our results do not dispute these findings, but also reveal a large diversity in the NO_2_^−^ utilization metabolisms in the ODZ and the taxa performing them. Based upon MAG relative abundance and normalized marker gene abundance within metagenomes, we resolve a picture of nitrogen cycling across the ODZ (Figure 1). We find a diversity of nitrogen metabolisms with the key intermediate NO_2_^−^ partitioned amongst anammox, DNRA, and denitrifying microbes in the ODZ, along with a large nitrifier contribution in the oxycline. The relative dominance of these metabolisms may be driven by competition for carbon along with bioavailable nitrogen, as well as enzyme tolerances for oxygen and interspecies interactions. The modularity of denitrification genes, and possibly other nitrogen cycling genes, may enable organisms to acquire these metabolisms as needed via mechanisms such as horizontal gene transfer. The possession of a gene does not necessitate its active transcription and function, and the regulation of denitrification genes, particularly in partial denitrifiers, remains to be fully comprehended. However, the broad patterns we find in gene content represent a metabolic potential present within microbes across ODZs and depths and reflect the adaptive processes shaping these communities.

Much remains to be discovered about nitrogen cycling within the ODZ and the communities performing these metabolisms. Importantly, integrating genomic information from MAGs with biogeochemical tracers and reaction rates for nitrogen cycling processes is a necessary step to identify the major microbial players in this system, resolve their activities in the water column, and predict how these communities will respond to and shape the global nitrogen cycle under changing climate conditions. Further studies on the full range of sequence space for given nitrogen cycle proteins requires a closer look into microbial physiology and metabolism in environmental microbes, possibly through culture efforts targeting understudied ODZ taxa.

## Data availability

MAGs will be submitted to NCBI. Metagenomic assemblies will be deposited to NCBI.

## Code availability

Code for metagenome assemblies, MAG generation, and bioinformatics analyses will be available on GitHub.

## Author contributions

IHZ and ARB conceptualized this study. IHZ assembled metagenomes and MAGs, conducted all bioinformatics analyses, and drafted the paper. AJ, XS, and BBW collected DNA samples, oxygen, and nutrient data and performed sequencing for metagenomes from this study. SGF and BBW provided methodological advice and guidance for the direction of this study. All authors provided feedback and proofread the paper.

