## supplemental figures and tables for "Genome-resolved metagenomics reveals abundant nitrate reducers and partitioning of nitrite usage within global oxygen deficient zones"

**Contents**

Supplementary Table S1

Supplementary Figures S1–S8

Supplementary Files S1–S3

| Reference | NCBI ID | Latitude (°) | Longitude (°) | Year | Ocean | Cruise & Station Number |
| --- | --- | --- | --- | --- | --- | --- |
| This study | N/A | 18.7 | -104.4 | 2016 | ETNP | RB-16-03  PS-S6 |
| This study | N/A | 17.2 | -110.7 | 2016 | ETNP | RB-16-03  14 |
| This study | N/A | 15.2 | 64 | 2007 | Arabian Sea | KNOX009  2N |
| This study | N/A | 16 | -105 | 2018 | ETNP | SR1805  PS2 |
| This study | N/A | 18 | -102 | 2018 | ETNP | SR1805  PS3 |
| Fuchsman et al. (2017) | PRJNA350692 | 17 | -106.5 | 2012 | ETNP | TN278  136 |
| Fuchsman et al. (2017) | PRJNA350692 | 16.5 | -107.1 | 2012 | ETNP | TN278  BB2 |
| Glass et al. (2015) | PRJNA254808 | 18.9 | -108.8 | 2013 | ETNP | NH-1315  2 |
| Glass et al. (2015) | PRJNA254808 | 18.9 | -106.3 | 2013 | ETNP | NH-1315  4 |
| Glass et al. (2015) | PRJNA254808 | 18.9 | -104.5 | 2013 | ETNP | NH-1315  6 |
| Glass et al. (2015) | PRJNA254808 | 18.8 | -104.7 | 2013 | ETNP | NH-1315  10 |
| Tsementzi et al. (2016) | PRJNA323946 | 18.5 | -104.5 | 2013 | ETNP | OMZoMBiE  6 |
| Stewart et al. (2012) | PRJNA68419 | -20.1 | -70.4 | 2008 | ETSP | MOOMZ |
| Ganesh et al. (2014) | PRJNA217777 | -20 | -70.8 | 2010 | ETSP | BiG RAPA |

**Supplementary Table S1.** References, NCBI BioProject IDs, sampling latitude and longitude, sampling year, sampling ocean location, cruise name, and sampling station ID for each set of metagenomes included in this study. A full list of depths and associated oxygen and nutrient profiles can be found in Supplementary File S2.

**Supplementary Figures**

­­­­
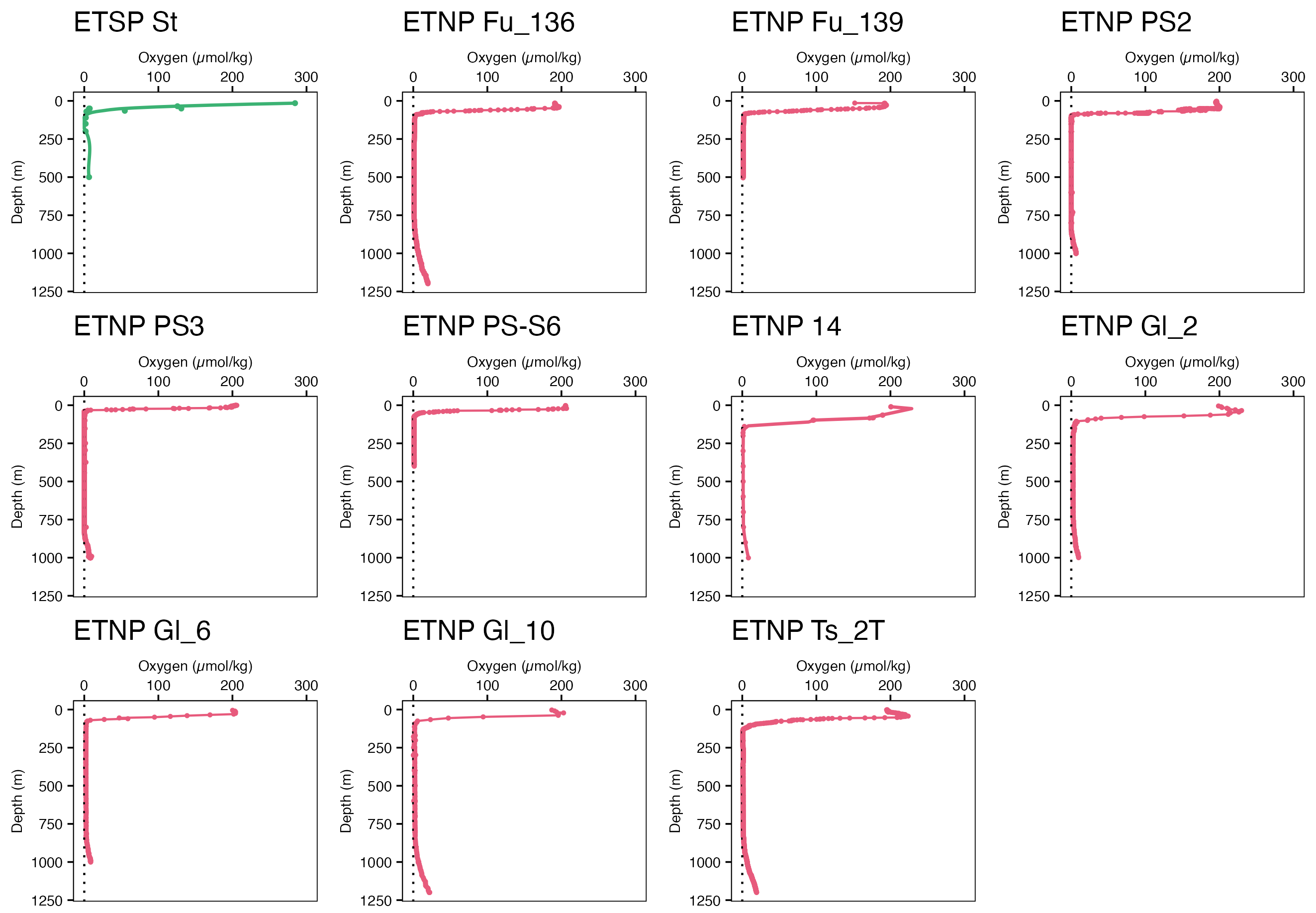

**Supplementary Figure S1.** Oxygen profiles from CTD casts for relevant sampling stations from which metagenomes were assembled. The naming scheme is as follows: St for Stewart profiles, Fu for Fuchsman profiles, Gl for Glass profiles and Ts for Tsementzi profiles, while letters and numbers following the underscore correspond to sampling sites following the original cruise naming scheme. Only 1 sampling site and profile corresponds to Stewart metagenomes. ETNP PS2, PS3, 14, and PS-S6 are profiles for metagenomes from this study. Three representative profiles are reported in Figure 2b.

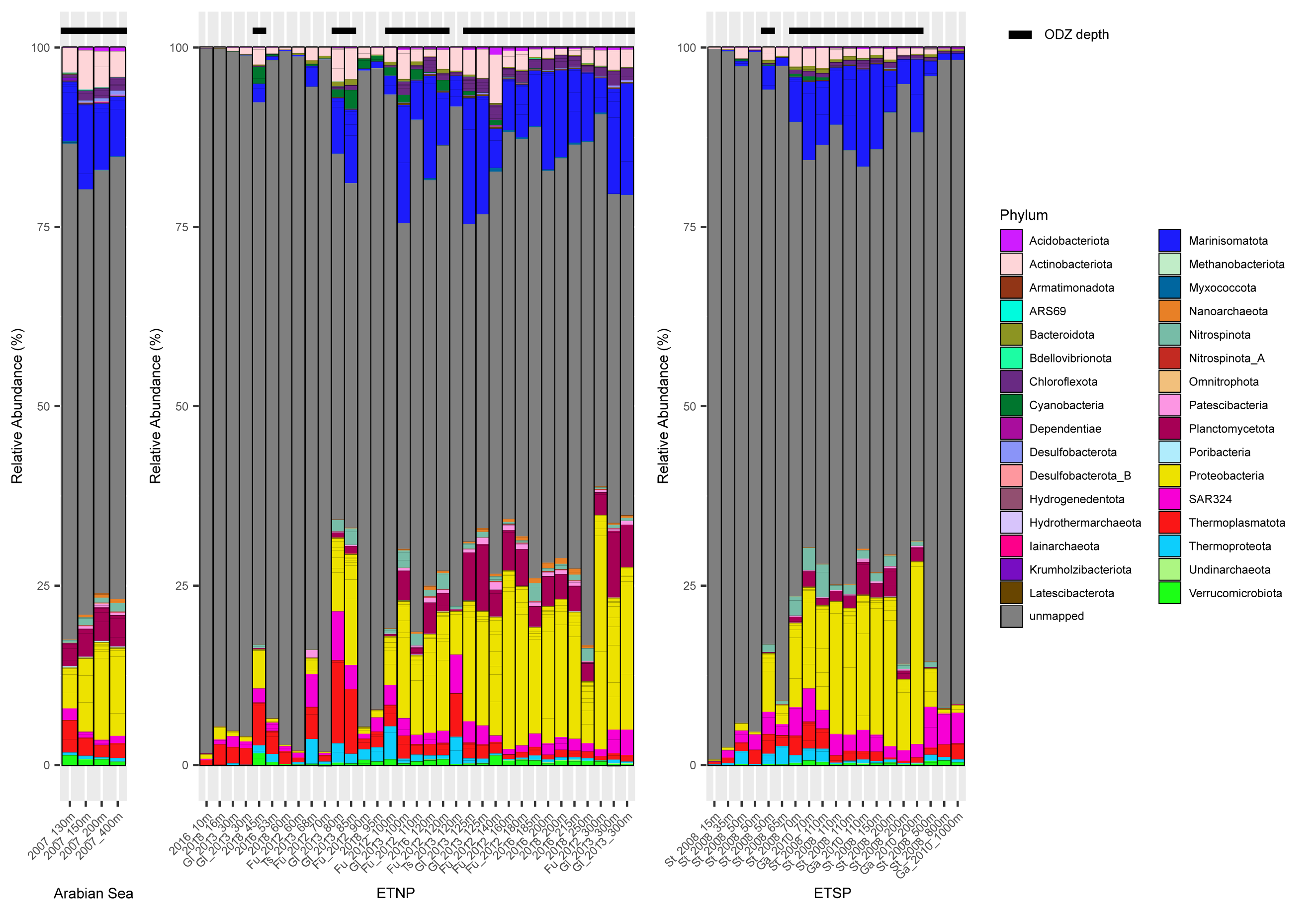

**Supplementary Figure S2.** Relative abundances of all dereplicated MAGs against all short reads from each metagenome, colored by phylum-level taxonomy. Grey bars indicate reads that were not mapped to any MAG in the collection. Metagenomes are arranged in the same order as in Figure 3a. Black bars above the graph indicate ODZ depths (O_2_ < 5 μM).

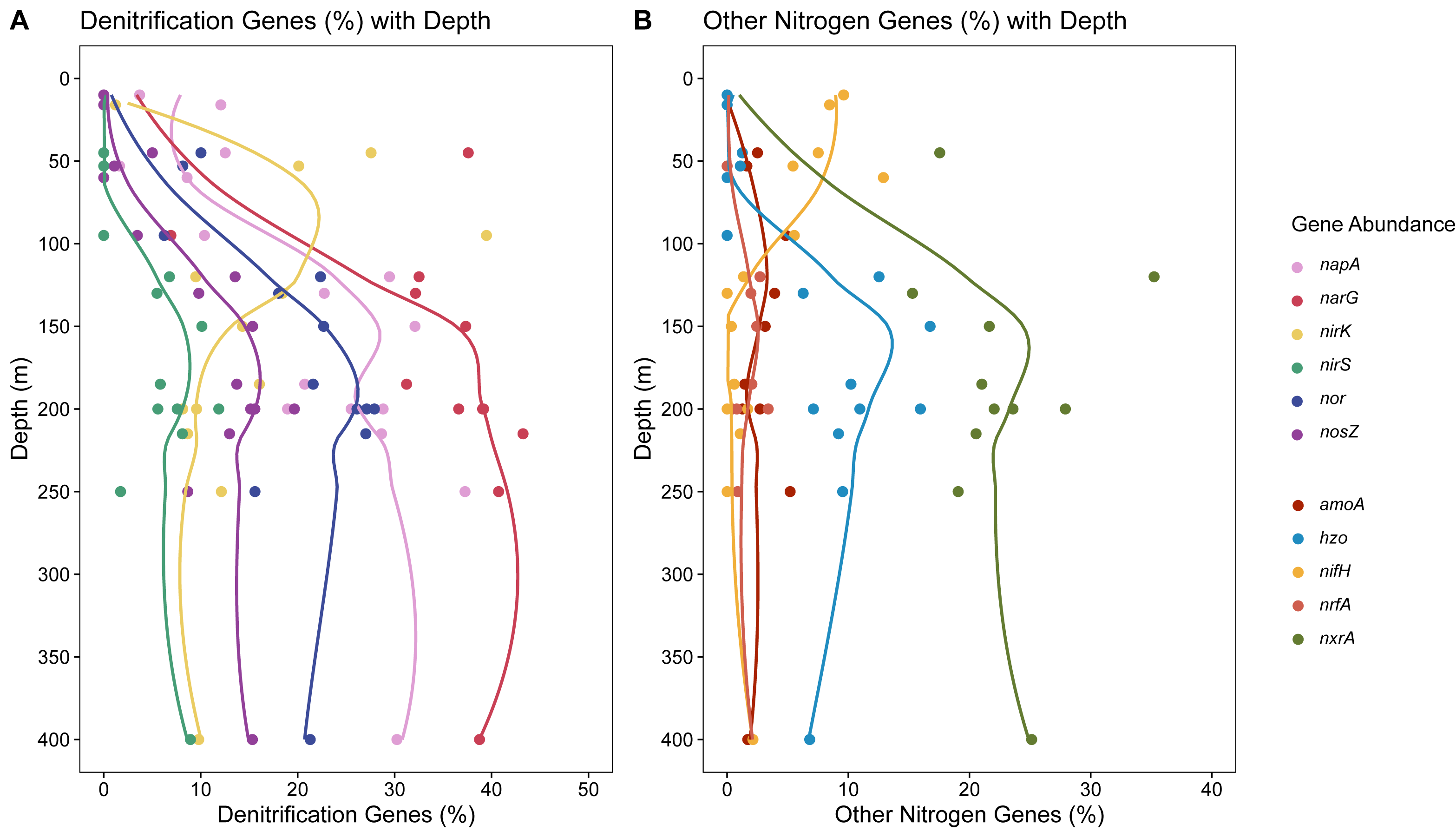

**Supplementary Figure S3.** a) Relative abundance of hits for each denitrification gene within individually assembled ETNP metagenomes from this study. Relative abundances are normalized against the average number of hits for single-copy genes *rpoB, rplB, gyrB, recA*, and *rpS3*. b) Relative abundance of hits for each non-denitrification nitrogen cycling gene within individually assembled ETNP metagenomes from this study, calculated as in a.

­­­­­­

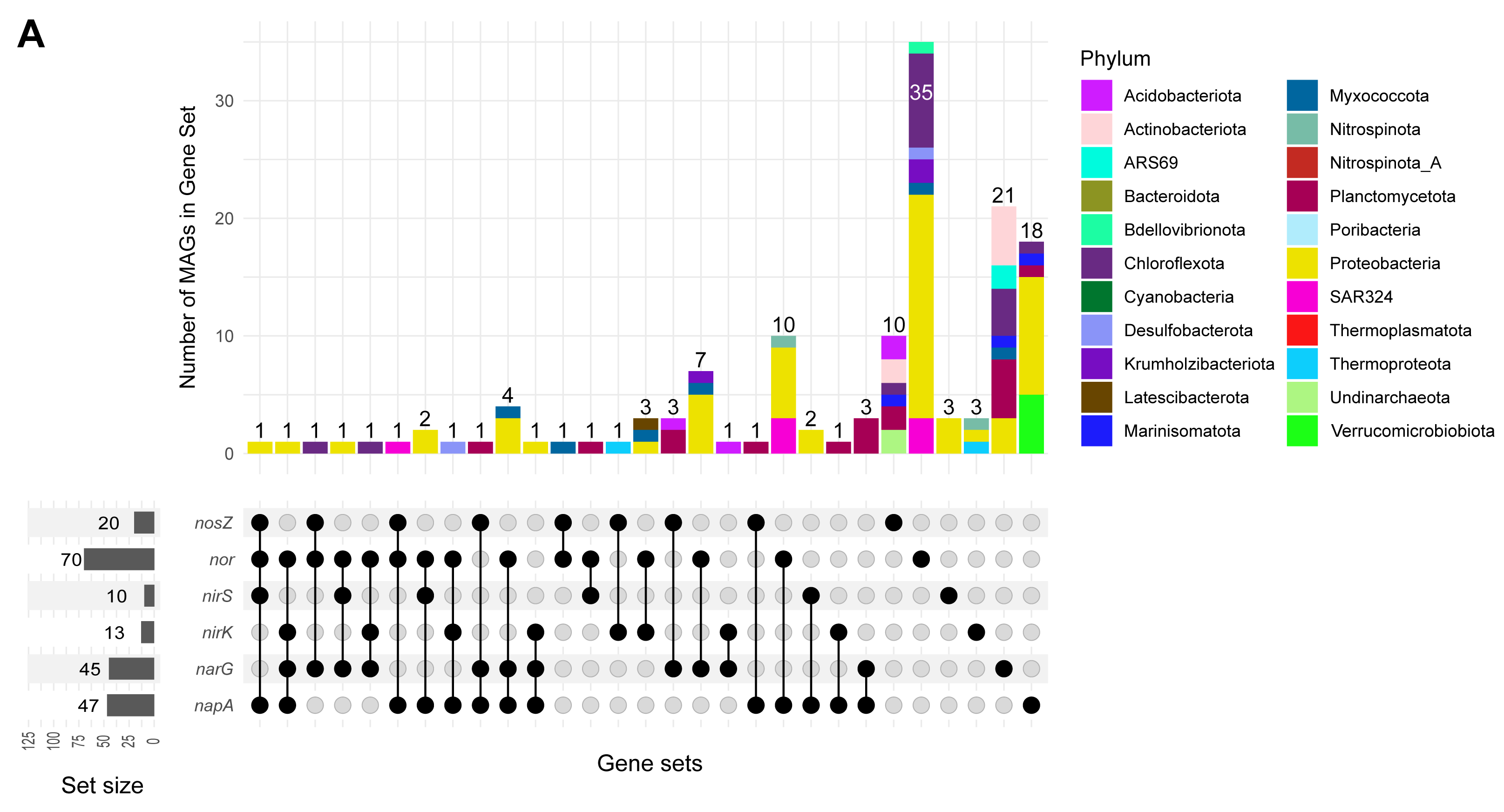

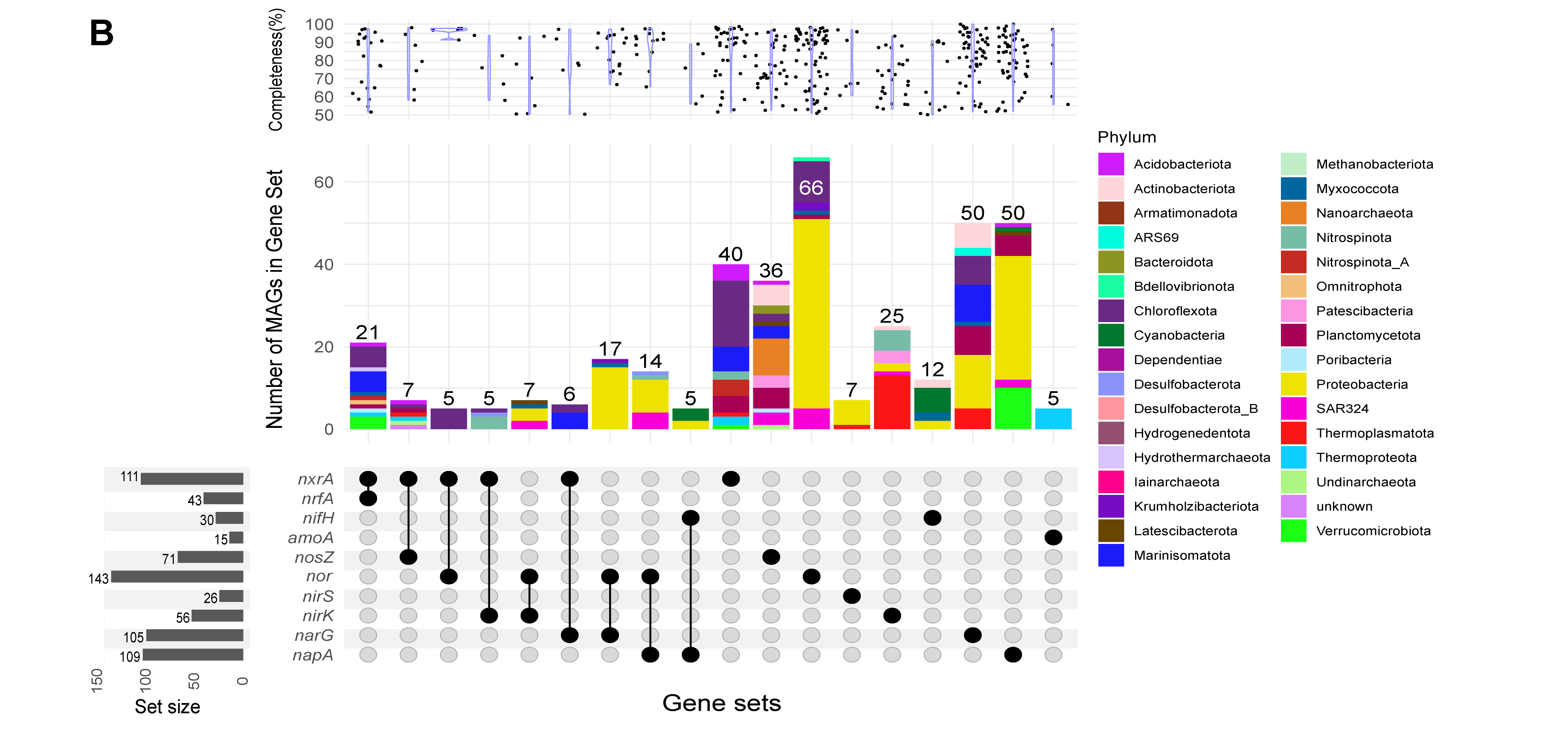

**Supplementary Figure S4.** a) The number of MAGs carrying each denitrification gene set within the ODZ MAG collection, showing only MAGs over 90% complete. Top panel shows the number of MAGs colored by phylum-level taxonomy, and bottom panel shows the genes within each gene set. Left bottom panel shows the number of MAGs carrying each specific denitrification gene. b) The number of MAGs carrying each nitrogen cycling gene set, including denitrification genes, within the ODZ MAG collection. Top panel shows the completeness (%) and the completeness distribution of the MAGs in that gene set, middle panel shows the number of MAGs colored by phylum-level taxonomy, and bottom panel shows the genes within each gene set. Left bottom panel shows the number of MAGs carrying each specific nitrogen cycling gene.

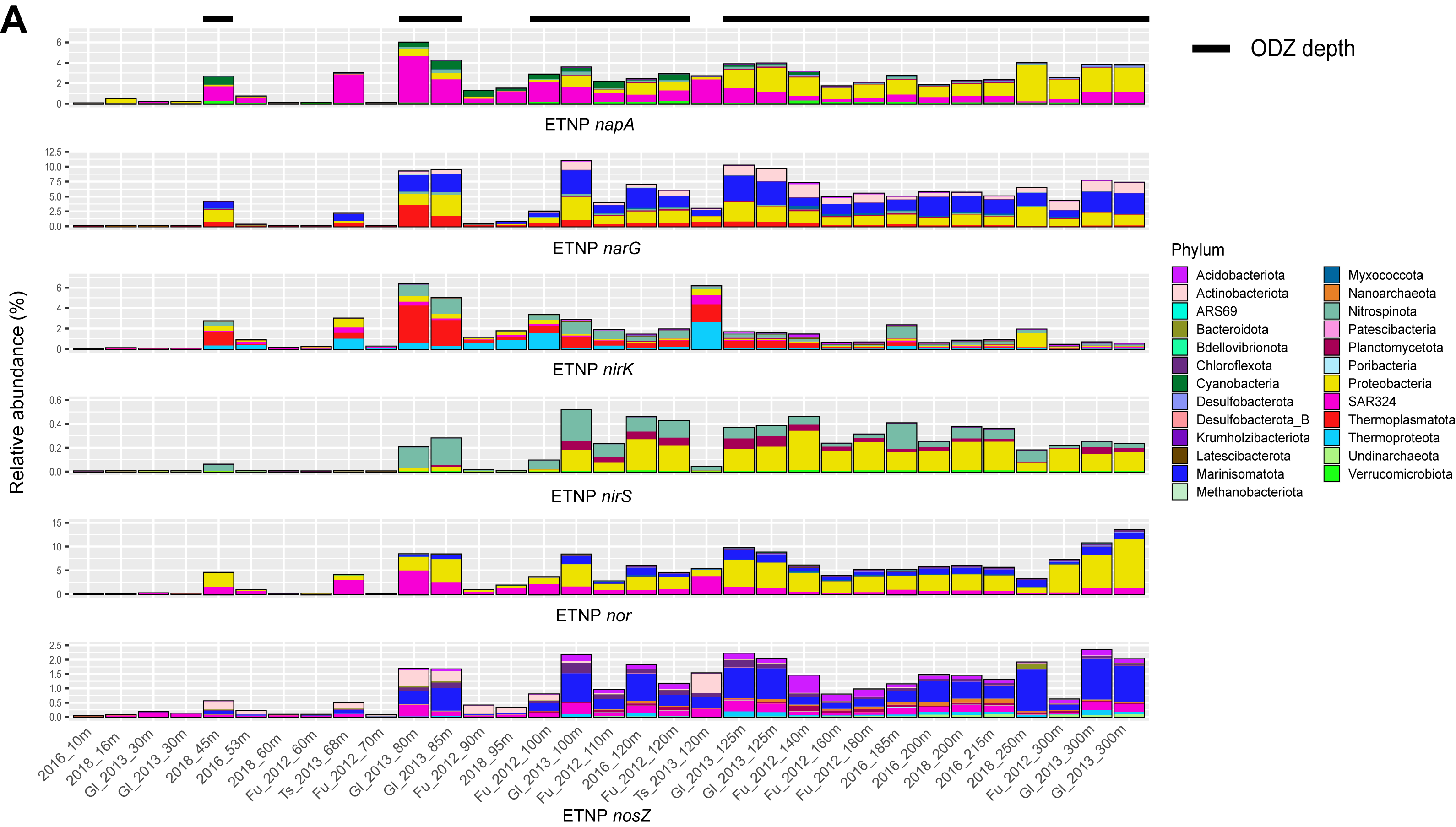

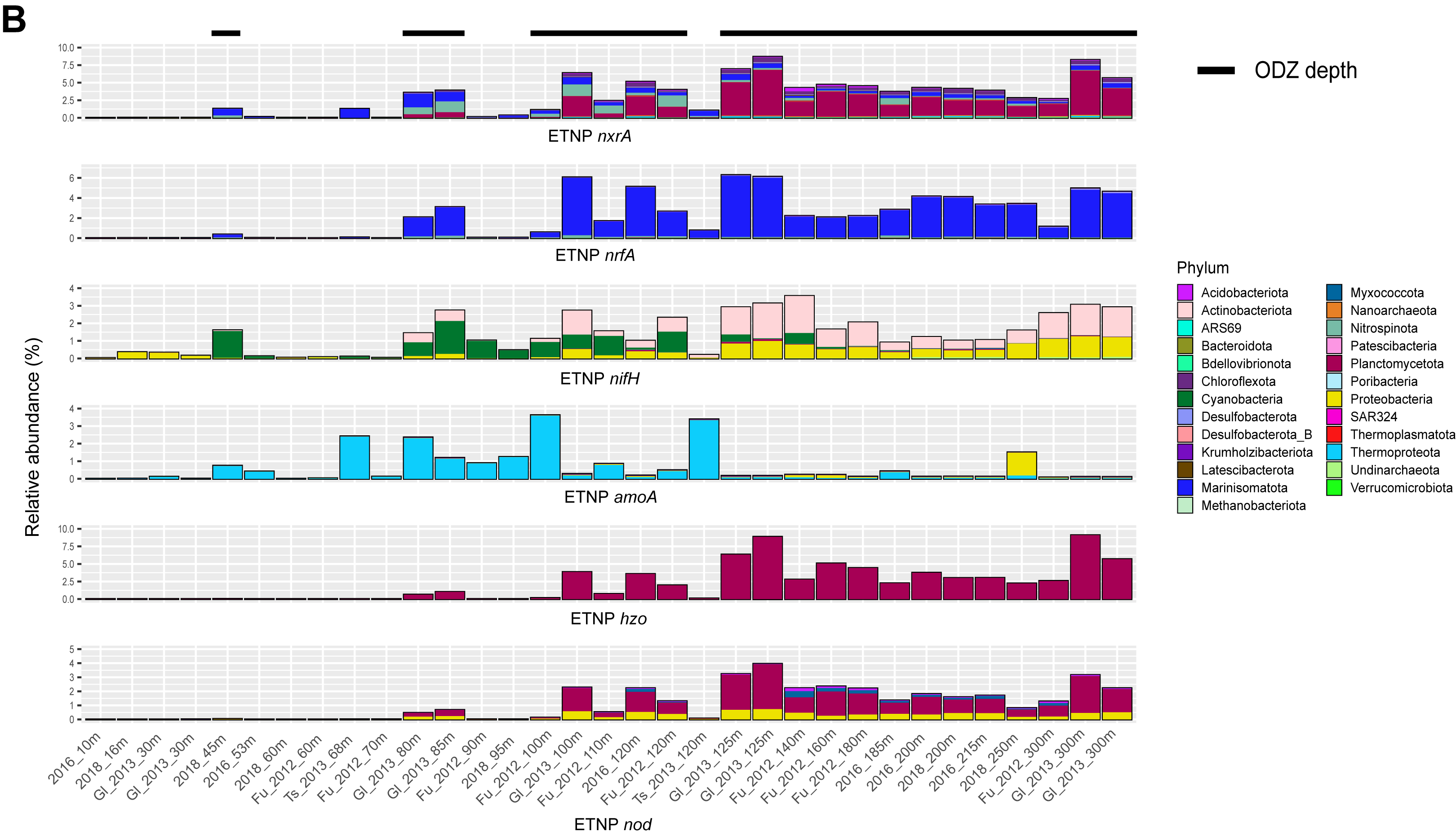

**Supplementary Figure S5.** Relative abundances of MAGs in the ETNP across all metagenomes carrying each a) denitrification gene b) other nitrogen-cycling genes. Black bars above the graph indicate ODZ depths (O_2_ < 5 μM).

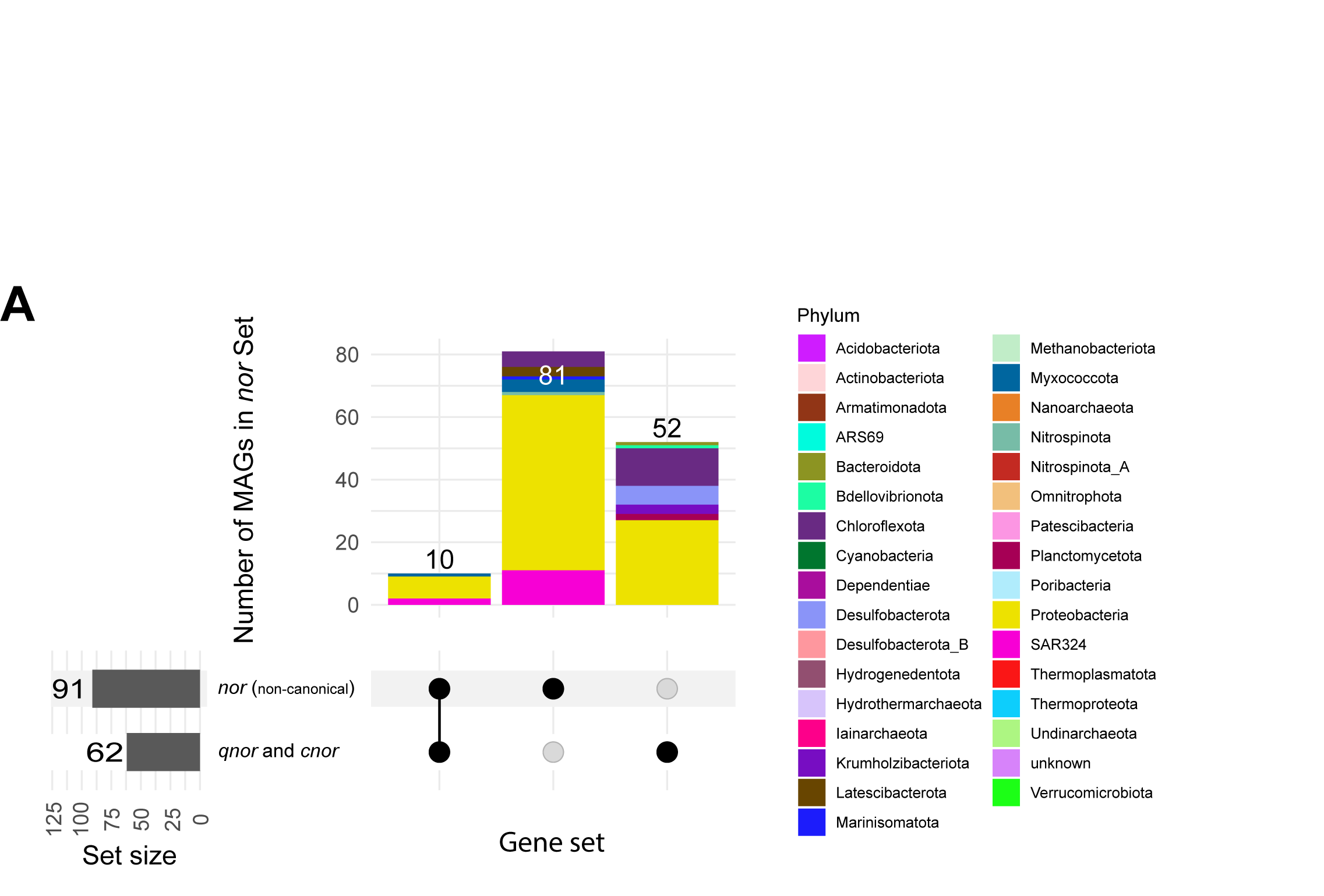

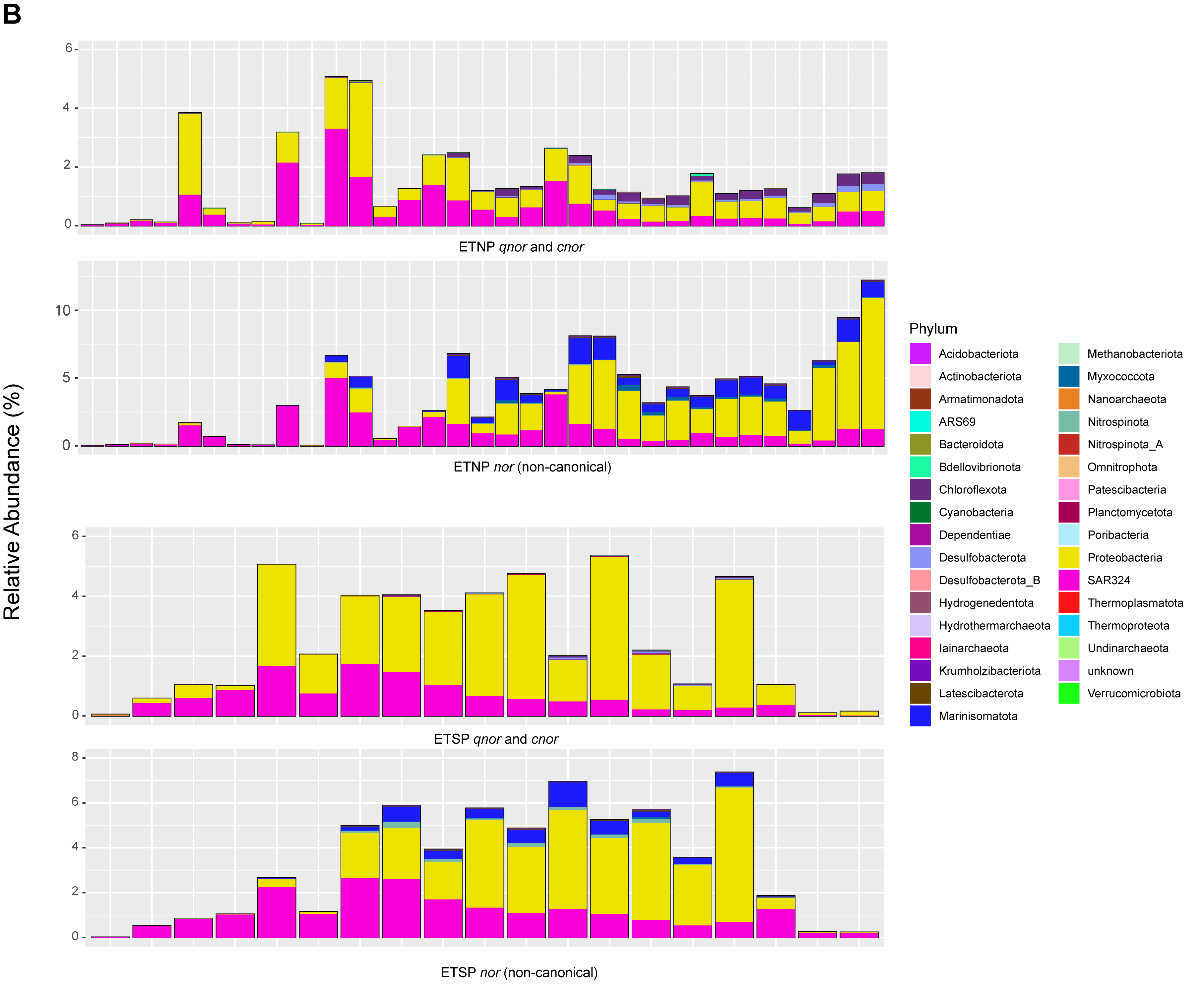

**Supplementary Figure S6.** a) The number of MAGs carrying canonical vs. non-canonical *nor* within the ODZ MAG collection. Top panel shows the number of MAGs colored by phylum-level taxonomy, and bottom panel shows the genes within each gene set. Left bottom panel shows the number of MAGs carrying each type of *nor*. b) Relative abundances of MAGs across all metagenomes carrying each type of *nor,* colored by phylum-level taxonomy. Top two panels correspond to ETNP, while bottom two panels correspond to ETSP.

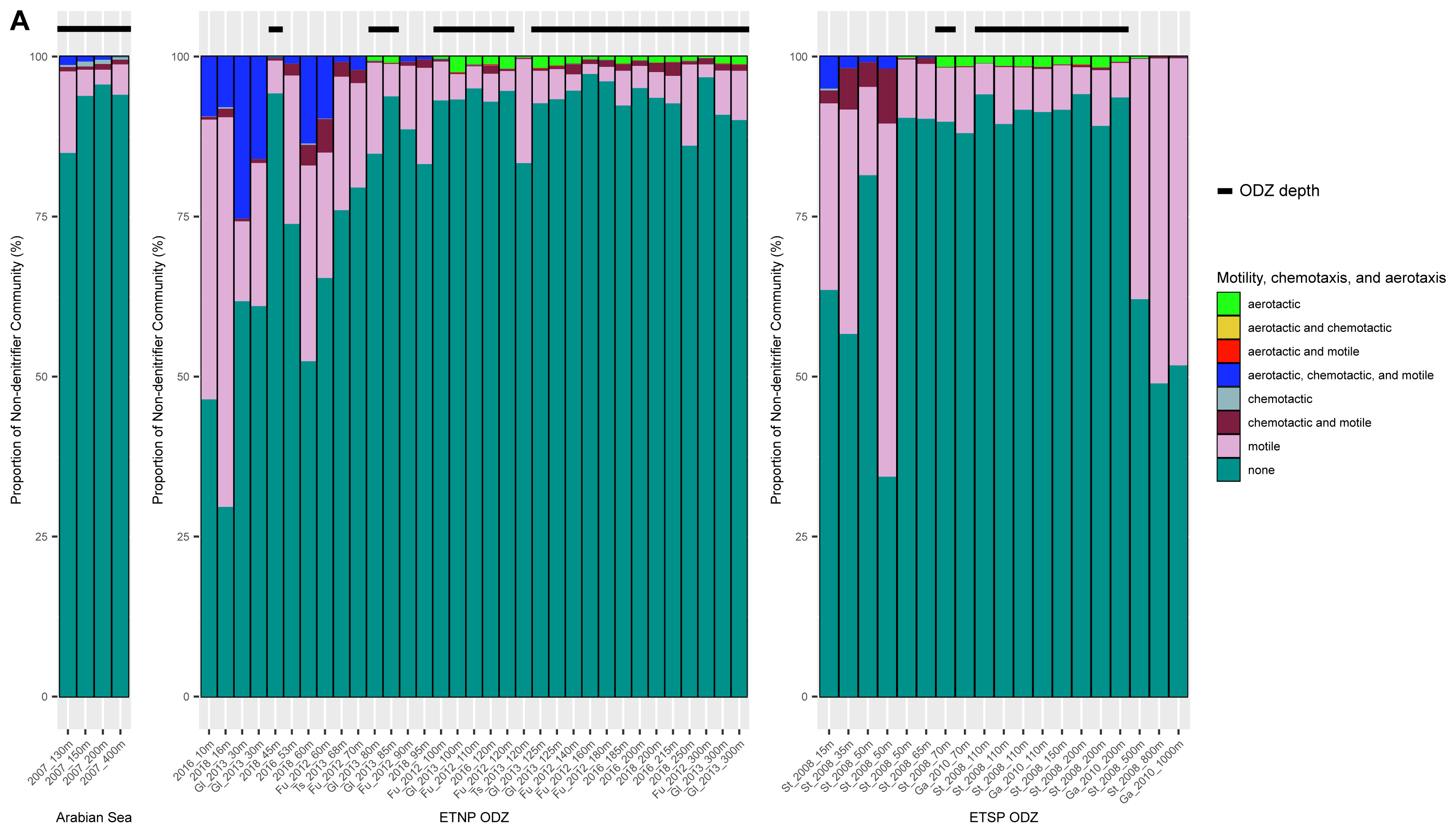

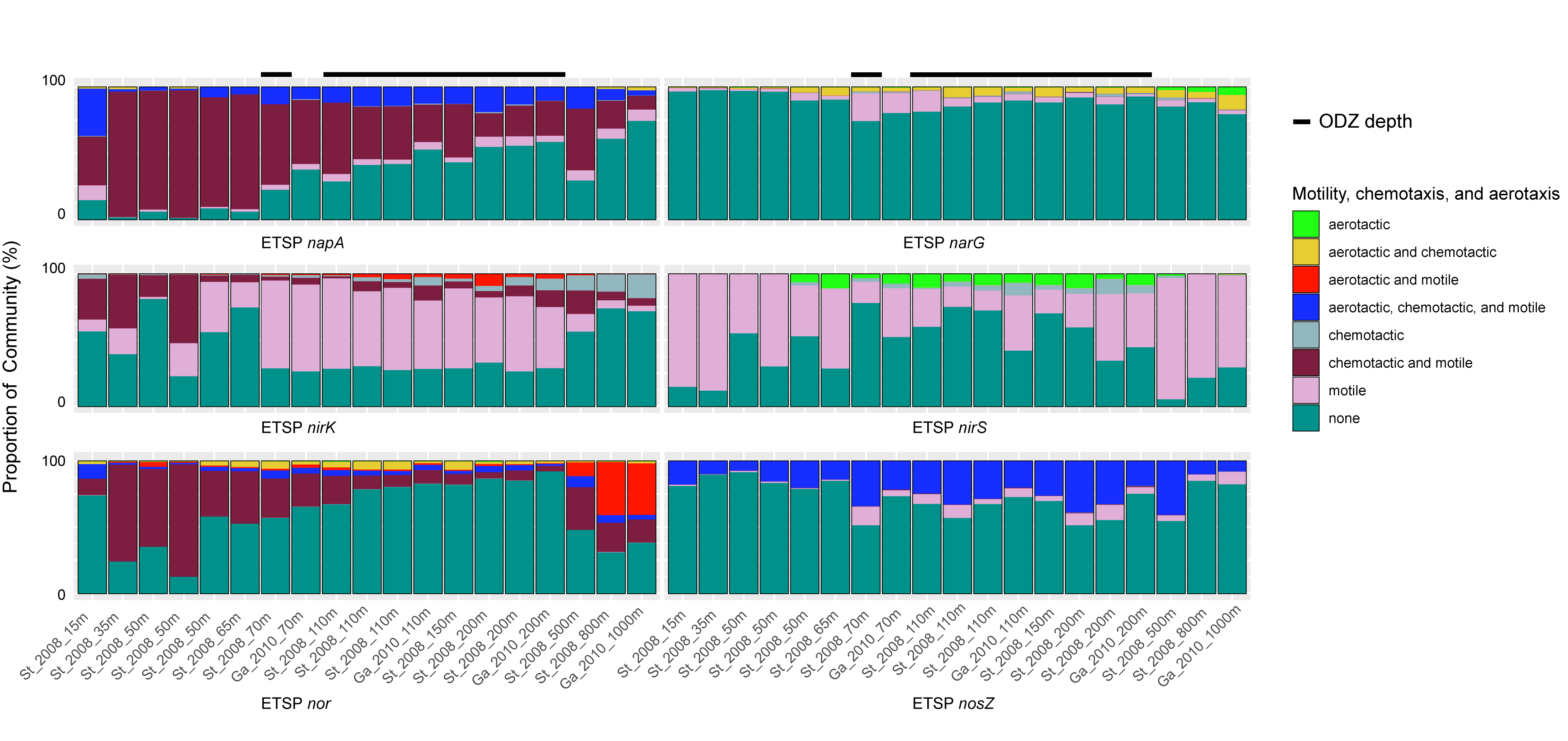

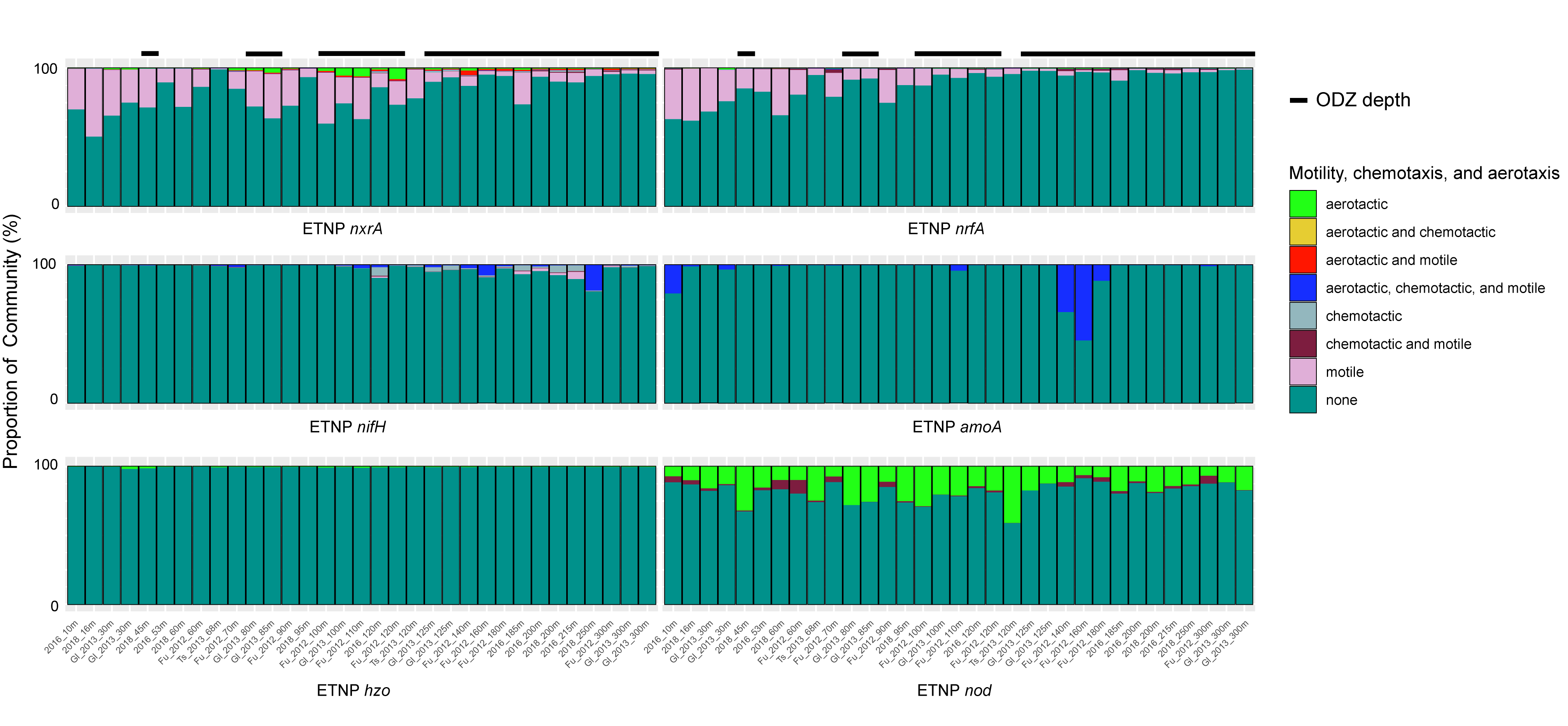

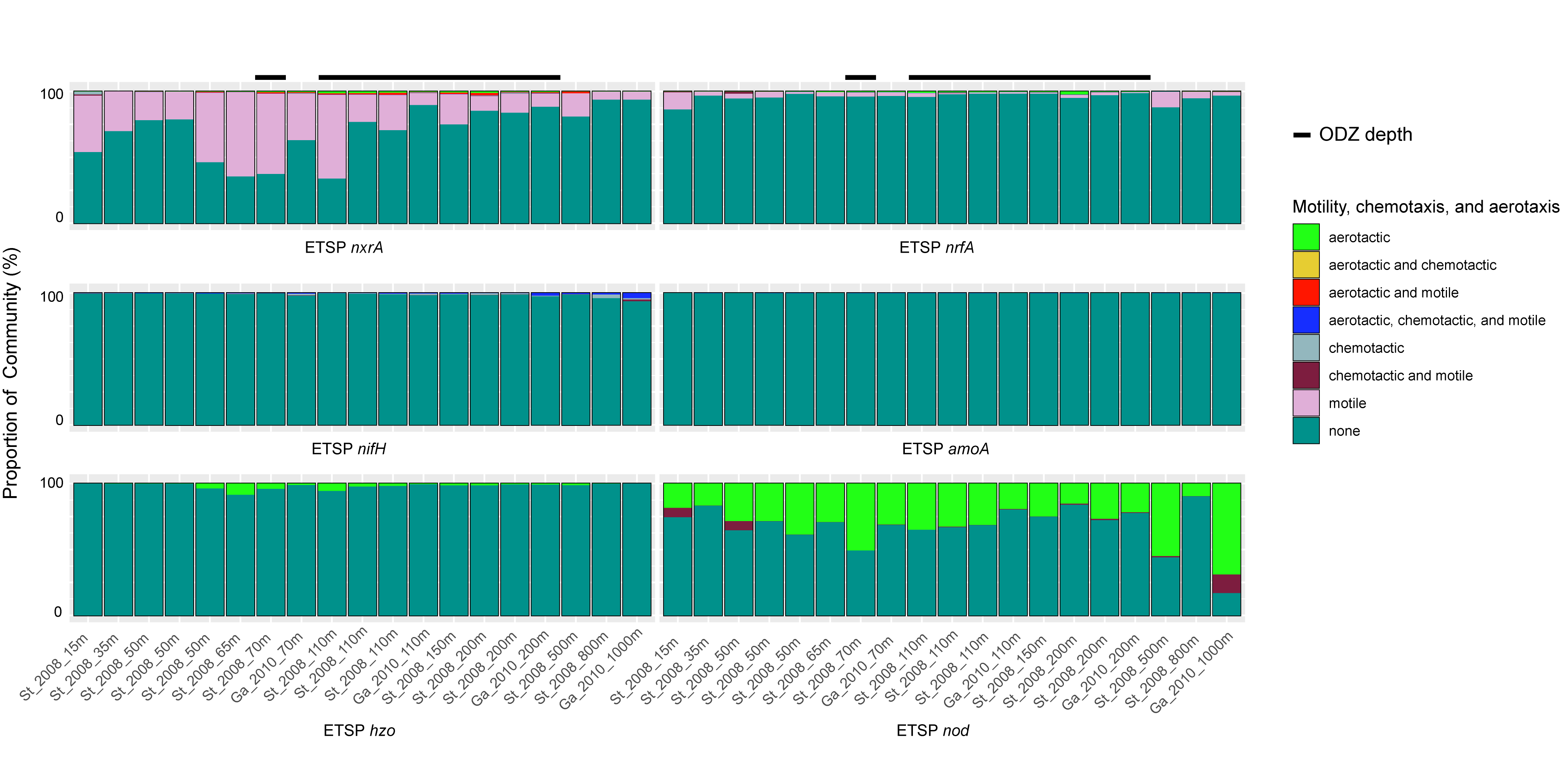

**Supplementary Figure S7.** a) Proportion of the non-denitrifying MAG community, scaled by relative abundance and colored by presence of motility, chemotaxis, and aerotaxis genes across all 3 ODZs, Graphs for the denitrifying community can be found in Figure 6a. b) Proportions of MAG communities carrying each queried denitrification gene, scaled by relative abundance and colored by presence of motility, chemotaxis, and aerotaxis, across all ETSP metagenomes. Graphs representing ETNP metagenomes can be found in Figure 6b. c, d) Proportions of the MAG communities carrying other nitrogen cycling genes, scaled by relative abundance and colored by presence of motility, chemotaxis, and aerotaxis for c) the ETNP and d) the ETSP. Proportions are calculated by (y /x)*100, where y = the relative abundances of all MAGs carrying the queried gene and falling into a specific motility, chemotaxis, or aerotaxis category and x = the relative abundances of all MAG carrying the queried gene.

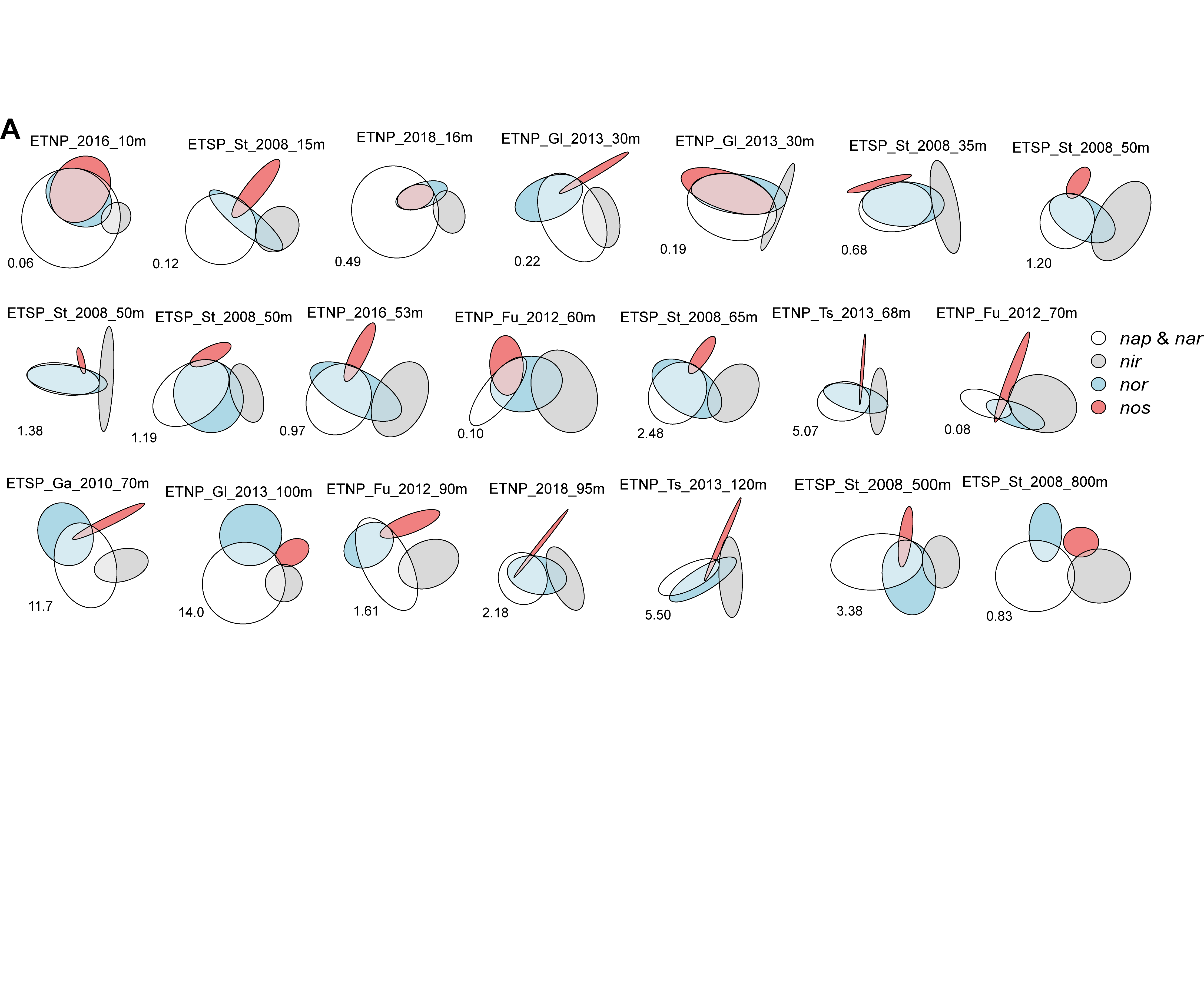

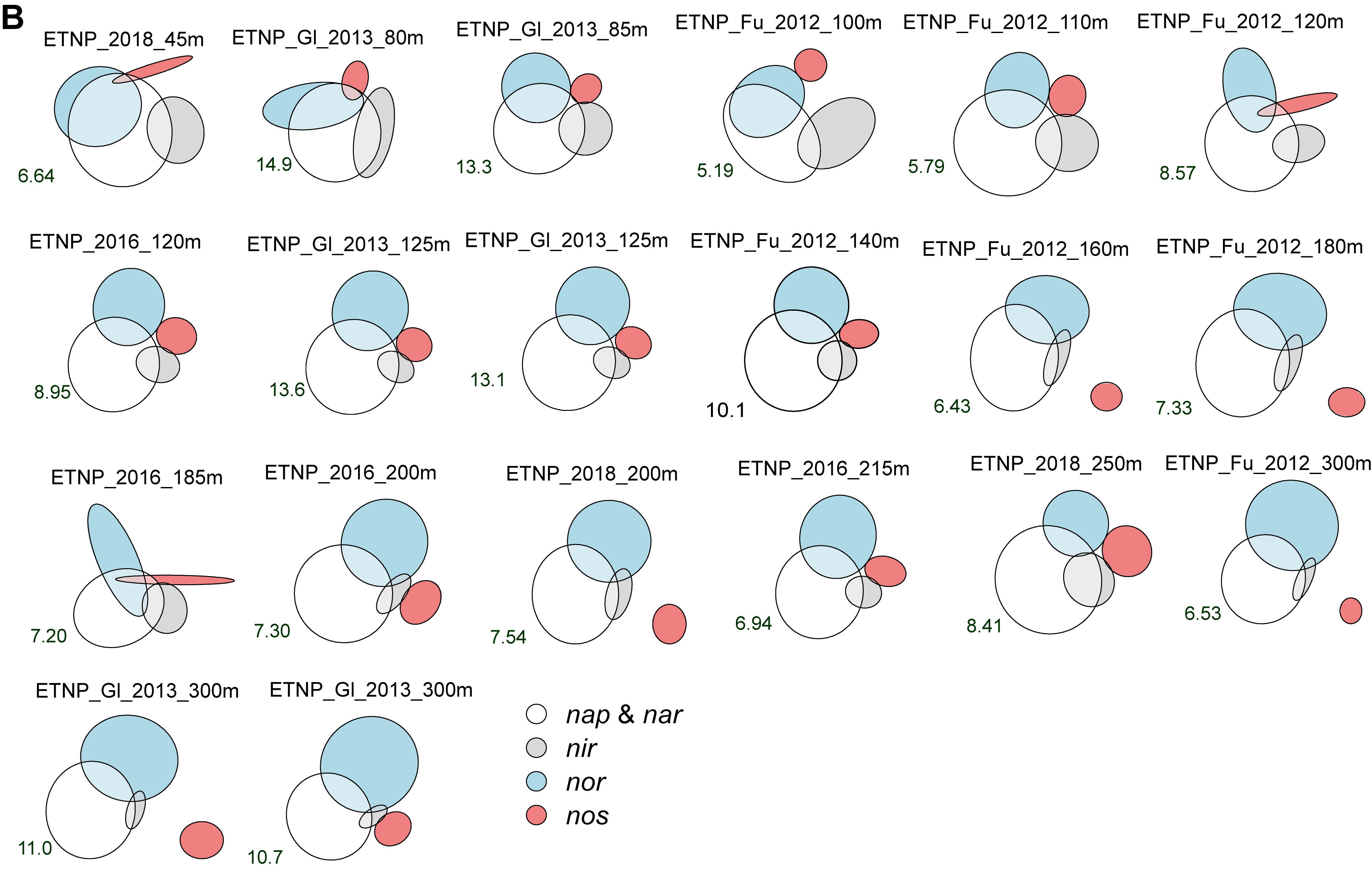

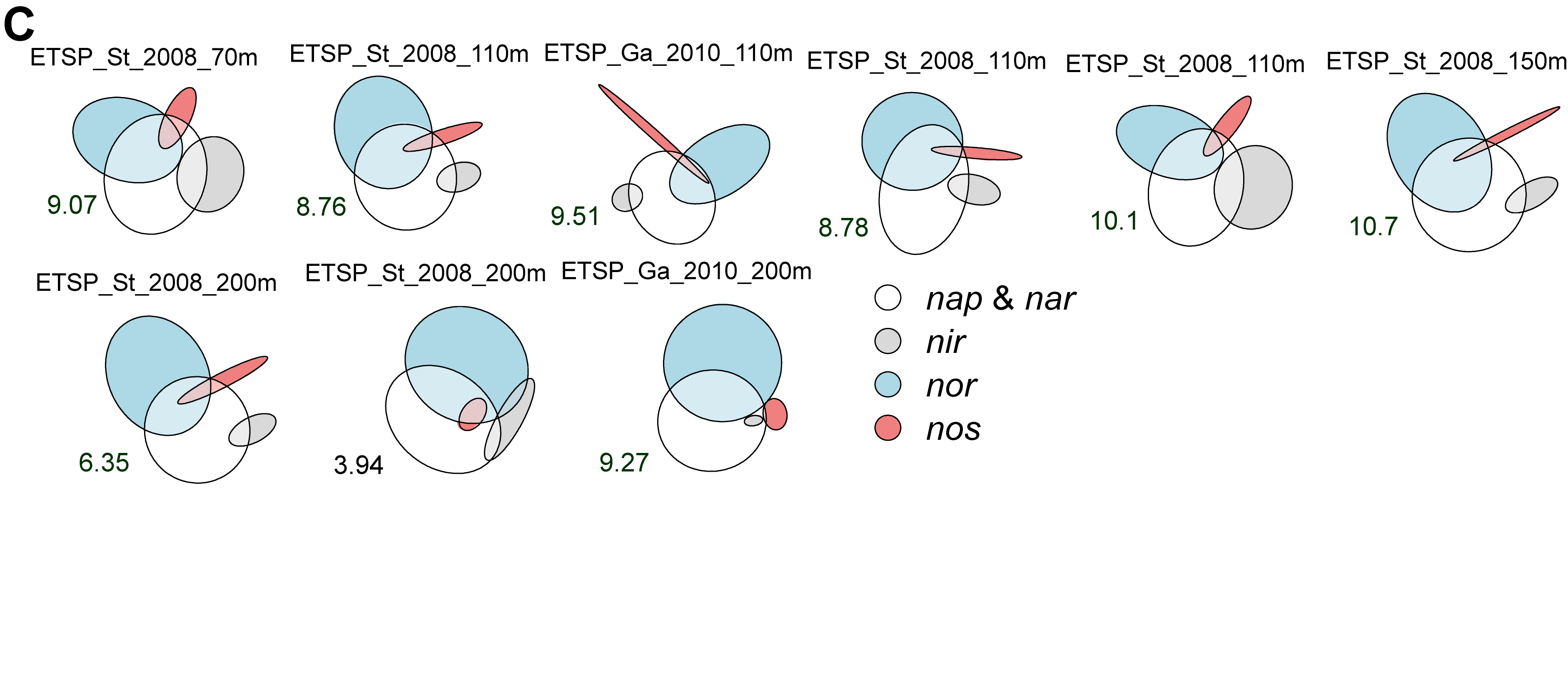

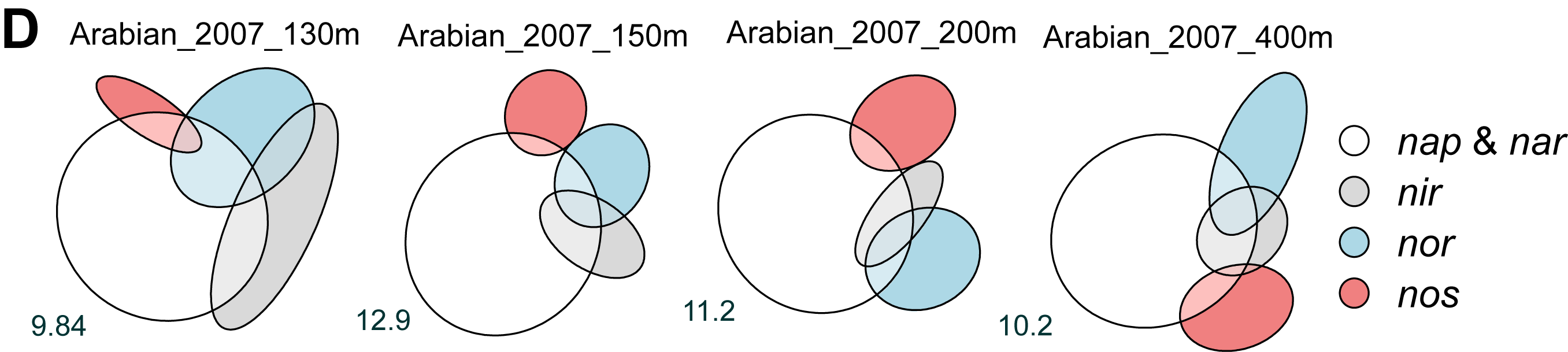

**Supplementary Figure S8.** Euler diagrams for a) non-ODZ depths from the ETNP and ETSP b) ETNP ODZ depths c) ETSP ODZ depths d) Arabian Sea ODZ depths. Circles and intersections are scaled to the total relative abundance of all MAGs possessing the genes for that step or step combination for each metagenome. The white circle corresponds to the relative abundance of MAGs with *napA, narG*, or both, and the numerical value for this relative abundance (%) is displayed on the left lower side of each diagram.

**Supplementary File S1.** All ODZ MAGs used in this study, with taxonomy, quality information, and gene presence/absence.

**Supplementary File S2.** All ODZ metagenomes included in this study, with associated oxygen data from CTD casts and nutrient data.

**Supplementary File S3.** Mapping results for all dereplicated ODZ MAGs against all metagenomes.
